# Structural Activation of DNA Unwinding by MCM8/9/HROB

**DOI:** 10.64898/2026.04.28.721408

**Authors:** Chuxuan Li, Colin To, Temitope M. Adeleke, David R. McKinzey, Yang Gao, Michael A. Trakselis

## Abstract

The minichromosomal maintenance MCM8 and MCM9 proteins form a heterohexameric complex that acts to unwind or remodel duplex DNA in DNA recombination and repair pathways. Mutations or absence of MCM8/9 have been linked to infertility, sex-specific deficiencies, and several cancers. Recently, HROB has been identified as a critical cofactor of MCM8/9; however, the mechanism underlying activation of MCM8/9 DNA binding and unwinding remain unclear. Here, we present dynamic structures of MCM8/9 with DNA, HROB and ATP analogs using cryo-electron microscopy. DNA binding induces a pronounced rotational rearrangement between the N-terminal DNA binding and C-terminal AAA^+^ ATPase domains, reorganizing DNA-binding loops into a staircase configuration that supports DNA engagement. Remarkably, HROB associates with both halves of the heterohexamer and drives a similar rotation prior DNA binding for localizing MCM8/9 to sites of crosslink damage and unwinding, culminating in a unified mechanistic model for MCM8/9 helicase function and its activation by HROB.

## Introduction

DNA helicases are prominent enzyme complexes that participate in DNA replication and repair by unwinding or remodeling duplex DNA. The minichromosomal maintenance (MCM) family of proteins is contained within the ATPases Associated with diverse cellular Activities (AAA^+^) superfamily, utilizing ATP hydrolysis to propel translocation and unwinding^1^. Archaea usually encode one or more MCM proteins that each assemble into a homohexamer for unwinding genomic DNA during DNA replication^2^. Within the eukaryotic MCM family, there are two evolutionarily conserved heterohexameric helicase complexes: MCM2-7 and the paralog complex, MCM8/9^3^. MCM2-7 is known to interact with four GINS proteins (SLD5/PSF1/PSF2/PSF3) and the cell division cycle protein 45 (Cdc45) to form the primary replicative CDC45-GINS-MCM2-7 (CMG) helicase complex utilized for DNA unwinding in eukaryotes^4–6^. In contrast, the latently evolved and less characterized MCM8/9 helicase complex appears to be integrated in multiple DNA damage mitigation strategies including homologous recombination (HR) and DNA fork protection, intersecting with the Fanconi anemia pathway^7–9^. Although the MCM helicases have differing cellular roles, the subunits share high sequence homology with multiple conserved motifs and structural domains^10^. MCM proteins are primarily composed of stacked C-terminal (CTD) and N-terminal domains (NTD)^8^. The NTD contains DNA binding loops and a zinc finger (ZF) motif for tight binding to DNA^11^. The CTD consists of the canonical AAA^+^ domains: the Walker A motif for nucleotide triphosphate (NTPs) binding, the Walker B motif for NTP hydrolysis, and two DNA binding loops for DNA translocation^12,13^. Although the MCM homologs share many similarities in their structures, MCM9 is unique in that it has a largely unstructured C-terminal extension (CTE) which has been shown to include specific motifs for nuclear localization and interactions with RAD51^14^.

Mutations or altered expression of MCM8 and MCM9 are linked to sex-specific pathologies and cancer, however, the connection between the associated pathologies and mutations or deficiencies in MCM8/9 are still unclear^15–24^. Mutations in MCM8 or MCM9 cause symptoms of early menopause, amenorrhea, and small or absent ovaries, leading to infertility characterized by primary ovarian failure or insufficiency (POF/POI), delays in puberty, and short statures^18–22,24^. Although less well studied, men are analogously afflicted with symptoms of azoospermia and small or absent testes^23^. MCM8/9 appears to not be required for DNA replication, but the genes are more highly expressed and active in S-phase^9,25^, and their absence delays cell cycle progression^8^. Altogether, the current evidence points toward a role for MCM8/9 in homologous or heterologous recombination during DNA replication or fork stalling in both mitotic or meiotic cells^7,8,14,26–29^. MCM8/9 interacts with FANCD_2_ during interstrand crosslink (ICL)^9^, the MRN complex during DNA resection^30^, and RAD51 for homologous recombination (HR)^14,31^. MCM9 was also found to interact with the mismatch repair (MMR) proteins (RFC, MSH2, MSH3, MLH1, and PMS1), likely utilizing its unwinding activity to remove the mismatched strand^32^. Recently, the Homologous Recombination OB-Fold (HROB) protein (formally MCM8IP) was identified as an HR associated protein through both proximity-dependent biotinylation and CRISPR-Cas9 screening and found to interact directly with MCM8/9^33,34^. Loss of HROB displays similar phenotypes as loss of MCM8/9 with increased ICL sensitivities, defective gametogenesis, and a decrease in DNA synthesis. HROB has been shown to directly activate MCM8/9 DNA unwinding and several interfacial residues at the NTD of MCM8 and MCM9 subunits were identified with an AlphaFold model of the MCM8/9/HROB complex^35^.

In this study, we present a series of high-resolution cryo-electron microscopy (cryo-EM) structures of human MCM8/9 helicase in complex with the non-hydrolyzable ATP analog (ATPγS), DNA, and HROB. These structures reveal the conserved hexameric architecture of MCM8/9 complex and their DNA-binding and ATPase site motifs. The DNA binding loops in MCM8/9 are organized in a staircase-like conformation, where only one subunit interface lacks both ATPγS binding and DNA engagement, suggesting a hand-over-hand translocation mechanism, akin to other hexameric helicases. Moreover, our structures elucidate unique and drastic conformational dynamics of MCM8/9 upon cofactor binding compared to our previous structure with only ADP bound. ATPγS binding induces contraction of the ATPase sites and the DNA binding channel.

Single-strand DNA (ssDNA) binding drives large scale rotation between the NTDs the CTDs of MCM8/9, which facilitates DNA binding and translocation of MCM8/9. HROB binding at a single MCM8/9 interface drives further NTD-CTD rotations, which prime the MCM8/9 complex for ssDNA engagement. Protein pulldowns and DNA unwinding quantitatively validate influential interfacial HROB residues that span the NTD (*site i*) and the CTD (*site ii*) tiers, both of which are important for MCM8/9 compaction for unwinding activation. Interestingly, although ATP is considered to be the preferred nucleoside triphosphates required for MCM8/9 unwinding activation, deoxyribonucleotides, dATP and dGTP, are also able to stimulate MCM8/9 unwinding similarly.

Finally, complementation assays in HROB knockout cells validate specific residues at *sites i and ii* important for crosslink damage induced MCM9 nuclear foci formation. Together, our findings provide a structural framework, biochemical and cellular foundation for understanding MCM8/9 activation and function in genome maintenance. Given the synthetic lethality associated with HROB loss, our structures offer a blueprint for rational targeting of the allosteric MCM8/9/HROB axis in cancer therapy.

## Results

### MCM8/9 C-terminal domains rotate upon binding to ssDNA

Previously, *Hs*MCM8/9^680^ was co-expressed and purified for structural analysis by cryo-EM^3^. The structure was captured using the Walker B mutants of both proteins incubated with ATP and hairpin fork single-stranded (ss) DNA. However, the resulting structure only captured MCM8/9^680^ with five hydrolyzed ADP molecules within the ATPase sites. To further investigate the mechanism of MCM8/9 DNA binding and translocation, MCM8/9^680^ was incubated with the non-hydrolyzable ATP analog ATPγS, and a forked hairpin DNA substrate (25 nt 3′ ssDNA overhang, 10 nt 5′-ssDNA flap, and an 8 base-pair (bp) double-stranded (ds) stem). Cryo-EM imaging and 2D classification revealed high-resolution features and diverse particle orientations of the MCM complexes (**Supplementary Fig. 1**). 3D reconstruction resolved two conformations: an MCM8/9^680^/ATPγS complex (MCM8/9^ATPγS^) at 3.51 Å and a DNA-bound MCM8/9/ATPγS/ssDNA complex (MCM8/9^ATPγS^–DNA) at 3.58 Å resolution (**Fig. 1 & Supplementary Fig. 1**). In both structures, most bulky residues and nucleotide ligands are well resolved. Similar to our previous MCM8/9 structures, MCM8/9 forms a heterohexameric ring composed of alternating MCM8 and MCM9 subunits (**Fig. 1A-B**).

**Figure 1.**
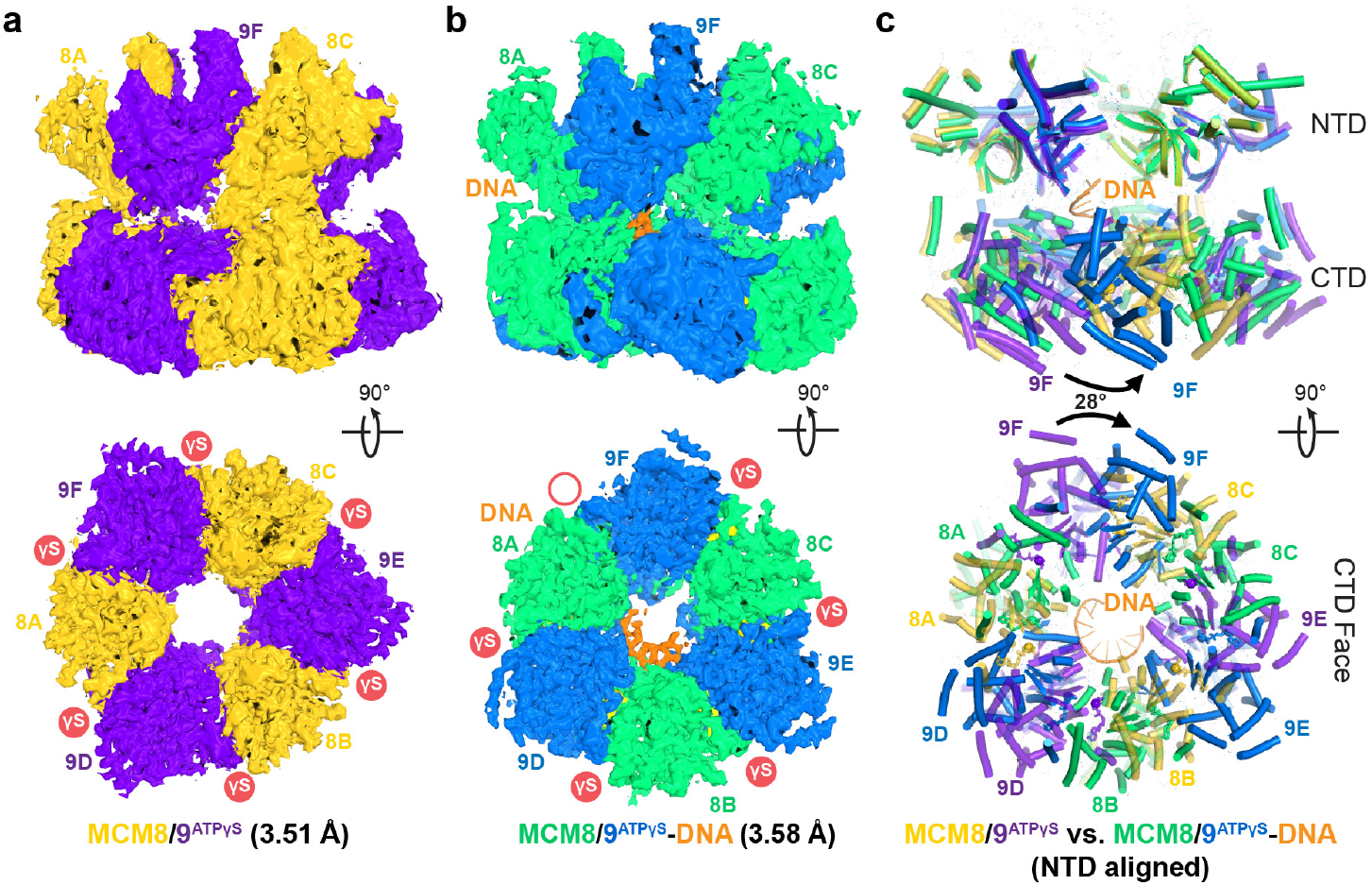
Cryo-EM structures of MCM8/9 complex with DNA. **a**, Overall cryo-EM density map of the MCM8/9ATPγS complex at 3.51 Å. Side view (top panel) and the bottom view from the CTD face of MCM8/9 (bottom panel) were shown. The MCM8 and MCM9 subunits were colored in gold and purple, respectively. The subunits were labeled as 8A, 9D, 8B, 9E, 8C and 9F clockwise in the bottom view of MCM8/9 hexamer. ATPγS occupancy is noted with closed red circle. **b**, Side view (top panel) and bottom view (bottom panel) of the MCM8/9ATPγS-DNA complex at 3.58 Å. The MCM8, MCM9, and DNA were colored in green, blue, and orange, respectively. **c**, Domain rotation upon DNA binding. In both the side view (top panel) and bottom view (bottom panel), the NTD rings from the MCM8/9ATPγS complex and the MCM8/9ATPγS-DNA complex were aligned. The movement of CTD ring was indicated by black arrows. The subunits were colored as in **a** and **b**. Protein loops were omitted for clarity.

The subunits were labeled 8A-9D-8B-9E-8C-9F in an anticlockwise direction around the MCM8/9 ring (**Fig. 1A**). Six ATPγS molecules are resolved in the MCM8/9^ATPγS^ complex, whereas only five ATPγS molecules are present in the MCM8/9^ATPγS^–DNA complex (**Supplementary Fig. 2A-B**).

Notably, in the MCM8/9^ATPγS^–DNA structure, a 9-nt ssDNA segment threads through the central channel of the MCM8/9 ring, shedding light on the working mechanism of MCM8/9 helicase (**Fig. 1B & Supplementary Fig. 2D**).

The MCM8/9 complex adopts a two-tiered architecture, comprising an NTD ring and a CTD AAA^+^ ring (**Fig. 1**). The MCM8/9^ATPγS^ complex is overall similar to our previously determined ADP-bound structure (MCM8/9^ADP^)^3^, with root-mean-square deviations (RMSDs) of 1.5 Å for the NTDs, 1.8 Å for the CTDs, and 1.7 Å overall. However, significant conformational changes occur upon DNA binding. When the NTDs are superimposed, the CTDs of both MCM8 and MCM9 subunits in the MCM8/9^ATPγS^-DNA complex undergo a 27-28° swinging rotation relative to their positions in the MCM8/9^ATPγS^ complex. This motion takes place through the flexible linker region between the two tiers (**Supplementary Fig. 3A-B**). As a result, the CTD ring rotates 28° clockwise in parallel relative to the NTD ring in the MCM8/9^ATPγS^-DNA complex (**Fig. 1C**). This rotation is possibly driven by the repositioning of DNA-binding loops in the MCM8/9^ATPγS^-DNA complex. In the absence of this movement, the DNA binding loop h2i would sterically clash with the NTD (**Fig. 2A**). Moreover, while the NTDs and CTDs are largely separated in the MCM8/9^ADP^ and MCM8/9^ATPγS^ structures (**Fig. 2B**), they establish inter-domain contacts in the MCM8/9^ATPγS^-DNA structure (**Fig. 2C**). Specifically, V546 in the MCM8 CTD engages in hydrophobic interaction with I131 in the MCM9 NTD, and E407 in the MCM9 CTD forms a hydrogen bond with the backbone of G305 in the MCM8 NTD.

**Figure 2.**
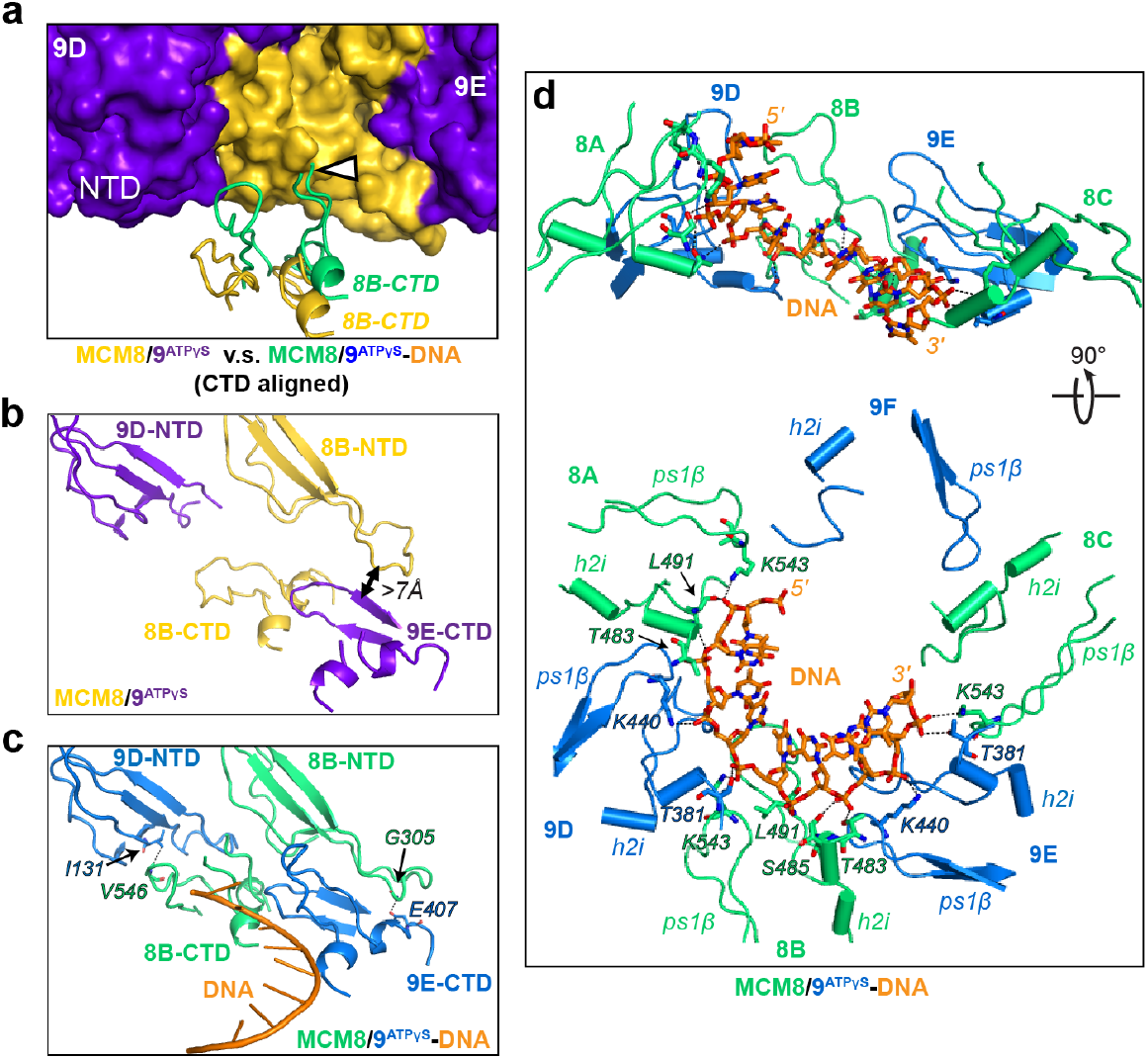
MCM8/9 interaction and conformational rotation with ssDNA. **a**, Potential steric clash between NTD and CTD upon DNA binding. The site of clash was indicated by a white arrow. The NTD rings of the MCM8/9^ATPγS^ complex (gold and purple) and the MCM8/9^ATPγS^-DNA (green and blue) complex was aligned. The NTD of the MCM8/9^ATPγS^ complex was shown as surface and the h2i loops from 8B subunits were shown as cartoon. NTD and CTD interactions in **b** the MCM8/9^ATPγS^ complex and **c** the MCM8/9^ATPγS^-DNA complex. The closest contact between 8B NTD and 9E CTD in the MCM8/9^ATPγS^ complex was indicated by a double-arrow, and the interactions between NTD and CTD in the MCM8/9^ATPγS^-DNA were indicated by the dotted lines, with the interacting residues labeled. **d**, Side view (top panel) and bottom view (bottom panel) of the DNA interaction loops in the MCM8/9^ATPγS^-DNA complex. The residues that interact with the DNA backbone were labeled.

In the MCM8/9^ATPγS^-DNA structure, the ssDNA is clearly resolved in the central channel of the CTD ring. The ssDNA adopts a helical conformation reminiscent of A-form nucleic acids, similar to that observed in archaeal MCM helicase as well as yeast and human CMG helicase structures (**Supplementary Fig. 3F-H**)^36–38^. The MCM8/9 complex interacts with the ssDNA via two conserved loops: the h2i and the ps1β (**Fig. 2D**). MCM8 residues L491 (backbone), T483, and K543, along with MCM9 residues K440 and T381, form a staircase-like conformation that contact the DNA backbone, with each subunit engaging approximately two nucleotides (**Fig. 2D**). Notably, the 9F subunit does not make any contact with the DNA, and no ATPγS density is observed at the 8A-9F interface (**Fig. 1B**).

### The OB-fold of HROB interacts at a single interface of MCM9

To characterize the MCM8/9 activation by HROB, we expressed and purified a core domain of HROB (residues 381-646, HROB^381-646^), which includes the OB-fold (residues 492-575) (**Supplementary Fig. 4**). HROB^381-646^ was incubated with MCM8/9^680^, ATPγS, and a hairpin fork DNA, and structures of the MCM8/9– HROB complex were determined in the presence of ATPγS and the forked DNA substrate. Similar to the MCM8/9^ATPγS^-DNA complex, cryo-EM data analysis produced two structures with distinct conformations: one corresponding to the MCM8/9^ATPγS^-HROB complex without DNA (MCM8/9^ATPγS^-HROB) at 4.16 Å and the second with bound DNA (MCM8/9^ATPγS^-HROB-DNA) at 2.96 Å (**Fig. 3 & Supplementary Fig. 5**). The MCM8/9 subunits in the MCM8/9^ATPγS^-HROB-DNA complex are almost superimposable with the MCM8/9^ATPγS^–DNA complex, with RMSDs of 0.7 Å, 0.6 Å, and 0.7 Å for NTDs, CTDs, and overall, respectively. In contrast, large-scale conformational changes and rotations are evident between the NTDs and CTDs in the MCM8/9^ATPγS^-HROB structure compared with MCM8/9^ATPγS^. When the NTDs are aligned, the CTDs of the MCM8/9^ATPγS^-HROB complex rotate ~25° clockwise relative to those in the MCM8/9^ATPγS^ structure, and ~4° anticlockwise relative to those in the MCM8/9^ATPγS^-DNA or the MCM8/9^ATPγS^-HROB-DNA complex (**Fig. 3C-D**).

**Figure 3.**
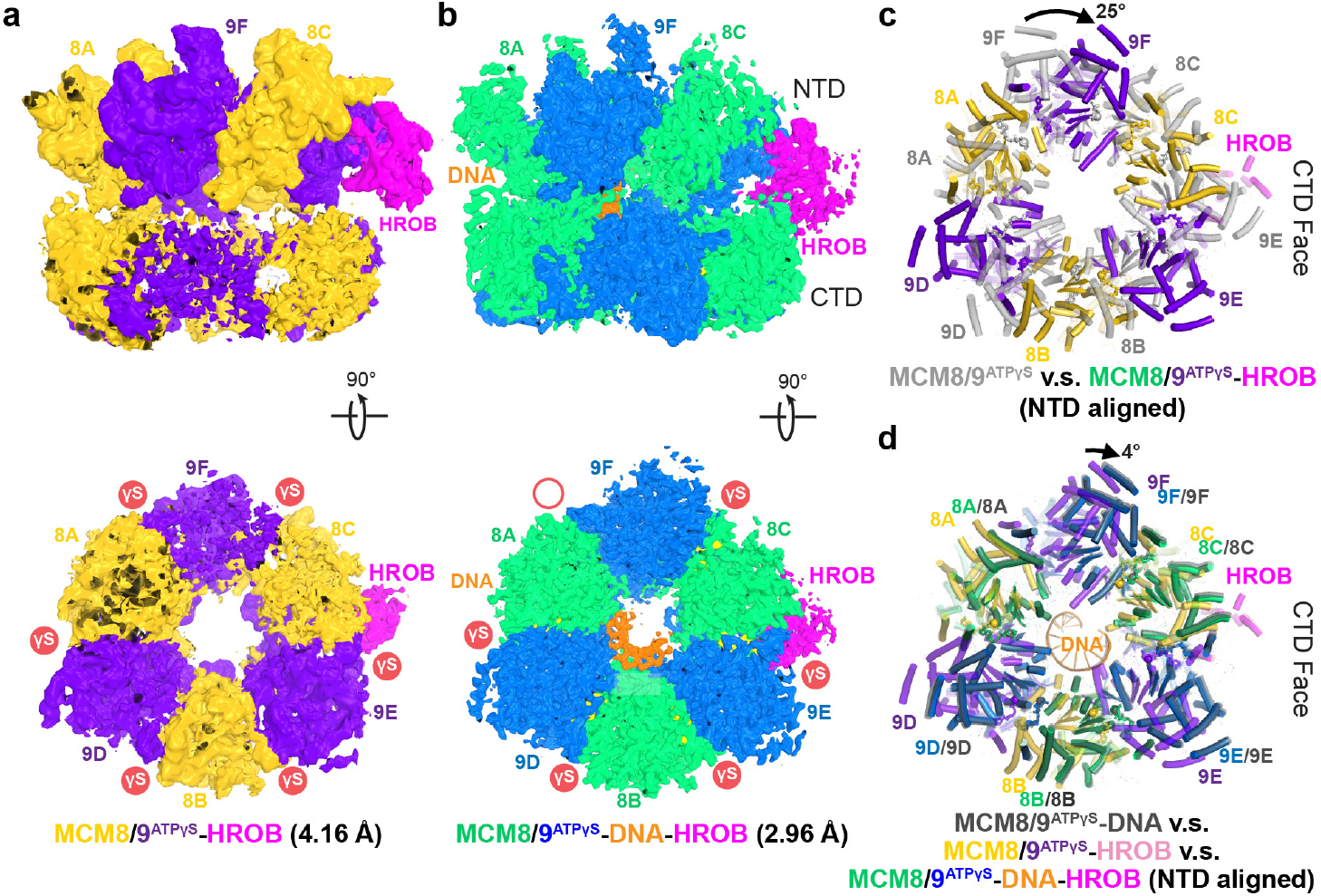
Cryo-EM structures of MCM8/9 complex with HROB. **a**, Side view (top panel) and bottom view (bottom panel) of the MCM8/9^ATPγS^-HROB complex at 4.16 Å. The MCM8, MCM9 and HROB subunits were colored in gold, purple, and magenta, respectively. The CTDs of 8A and 9F were flexible with poor density maps. ATPγS occupancy is noted with closed red circle. **b**, Side view (top panel) and bottom view (bottom panel) of the MCM8/9^ATPγS^-HROB-DNA complex at 2.96 Å. The MCM8, MCM9, HROB, and DNA were colored in green, blue, magenta, and orange, respectively. **c**, Domain rotation upon HROB binding. The NTD rings from different MCM8/9 complexes were aligned and only bottom views were displayed. In **c**, the MCM8/9^ATPγS^-HROB (gold and purple) complex was compared to the MCM8/9^ATPγS^ complex (light grey); whereas in **d**, the MCM8/9^ATPγS^-HROB (gold and purple) complex was aligned with the MCM8/9^ATPγS^-DNA complex (dark grey) and the MCM8/9^ATPγS^-HROB-DNA complex (green and blue). Protein loops were omitted for clarity.

In both MCM8/9^ATPγS^-HROB and MCM8/9^ATPγS^-HROB-DNA structures, cryo-EM densities for HROB can be clearly observed near the NTD of subunit 9E (**Figs. 3 & 4A-C**). In the overall moderate resolution MCM8/9^ATPγS^-HROB structure, the local NTD and HROB structures are better resolved, but the CTD is partially disordered, diminishing the overall resolution (**Supplementary Fig. 5**). In the higher resolution MCM8/9^ATPγS^-HROB-DNA structure, the NTD, CTD, and HROB are all well traced, including the residues at the 9E-HROB interface (**Fig. 4B**). Additionally, five ATPγS molecules and an extended stretch of 11-nt ssDNA are defined (**Fig. 3B & Supplementary Fig. 2C & E**). However, despite using a HROB construct spanning residues 381-646, only residues 475-576 could be refined (**Supplementary Fig. 2F**). Additional density beyond the OB-fold of HROB is observed adjacent to the CTD of 8C subunit, suggesting partial engagement of MCM8 with other HROB segments. To improve visualization of HROB, we performed local classification and refinement with a mask only covering HROB (**Supplementary Fig. 5A**). This yielded a more complete map for HROB, enabling modeling of additional segments corresponding to residues 395-407 and 441-475 for additional MCM interactions (**Fig. 4A-C & Supplementary Fig. 2G**).

**Figure 4.**
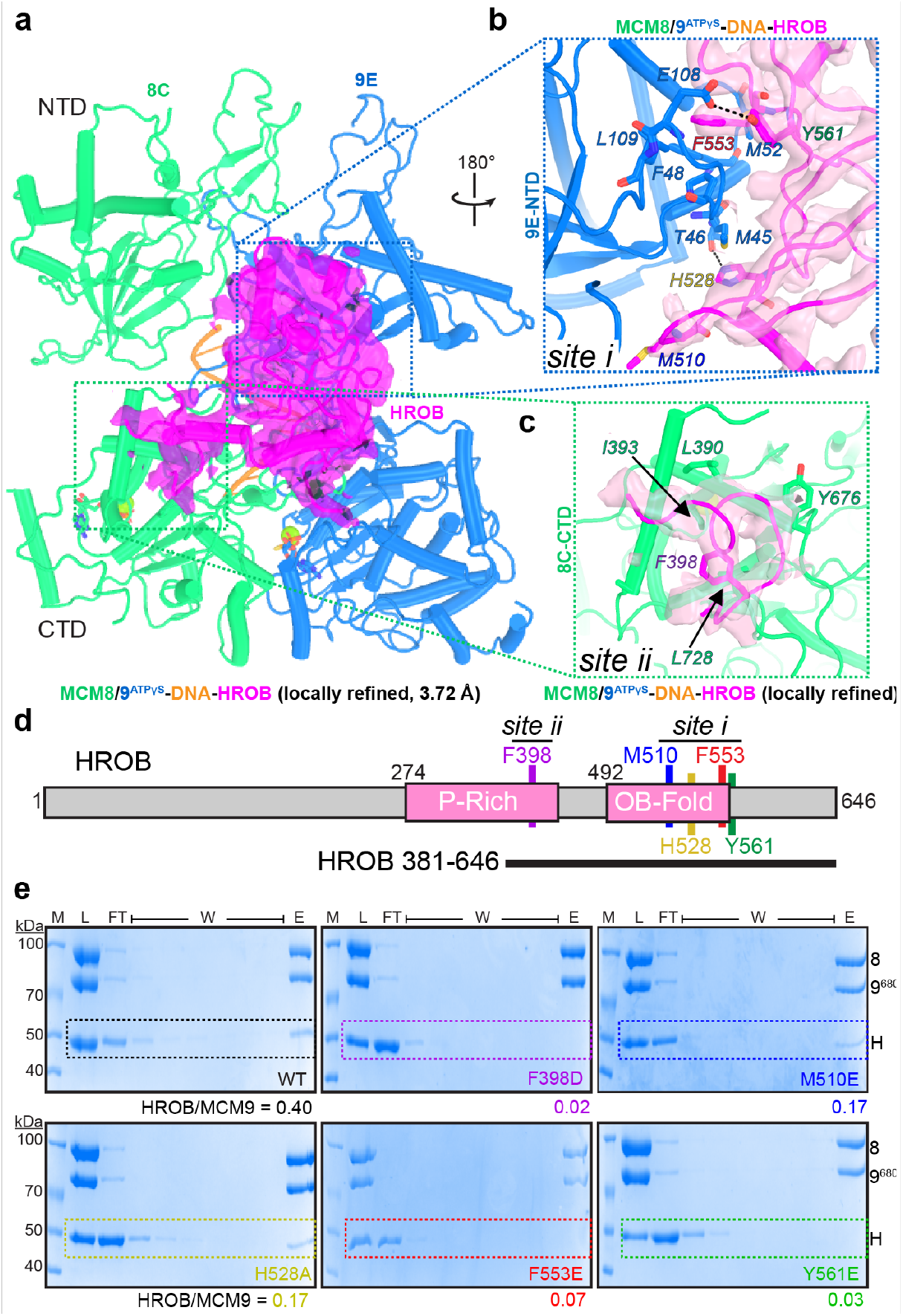
MCM8/9 interaction with HROB. **a**, HROB interaction with MCM8/9 in the locally refined MCM8/9^ATPγS^-HROB-DNA complex at 3.72 Å. The 8C and 9E dimer were shown as cartoon. The subunits were color-coded as in Figure 3 and the electron density for HROB was shown in magenta surface. The density map for HROB was derived from the MCM8/9^ATPγS^-HROB-DNA complex. Zoom-in view of HROB interaction with **b** NTD of the 9E subunit (*site i*) or the **c** CTD of the 8C subunit (*site ii*) are indicated as a dotted box. Sidechains at the interfaces were shown in sticks and the hydrogen-bonds were indicated by the dotted lines. **d**, Linear schematic of the HROB sequence showing the P-Rich and OB-fold domains in pink along with the targeted interaction residues to mutations indicated. Black bar indicates the HROB truncation construct used (381-646). **e**, Strep-Tactin XT magnetic beads used to immobilize Strep-tag-MCM8 to pull-down both MCM9 and HROB. Shown are representative gels that include markers (M), load (L), six washes (W), and the elution (E) with D-biotin for native HROB^381-646^ or site-specific mutations: F398D (purple), M510E (blue), H528A (mustard), F553E (red), and Y561E (green). The dashed colored boxes are highlighting the migration position for each HROB construct. The fractions below indicate the relative ratio of HROB to MCM9 in E.

### HROB interacts with MCM8/9 at two sites by inserting bulky, aromatic residues into hydrophobic pockets

Structural inspection reveals that HROB bridges the MCM8/9 NTD and CTD rings with two interfaces at the 9E NTD and 8C CTD (**Fig. 4A-C**). The 9E-HROB interface (*site i*) involves the interaction between the OB-fold domain of HROB and 9E NTD, which is extensive and buries approximately 981 Å^2^ of surface area. HROB residue F553 is inserted into a hydrophobic pocket formed by MCM9 residues F48, M52, P104, and L109 (**Fig. 4B**).

Additionally, HROB residues Y561 and H528 form hydrogen bonds with MCM9 residues E108 and T46, respectively, and M510 contacts the MCM9 at the bottom of the interface. The MCM9–HROB interface is similar as predicted in previous AlphaFold (AF) models^35^ (**Supplementary Fig. 6A-B**). The HROB subunit is with a RMSD of 2.2 Å between the AF-predicted model and our cryo-EM structures, with the portion closer to the MCM9-HROB interface better predicted, while the distal portion is less consistent. However, the subunit movement between the MCM8/9 NTD and CTD tiers was not predicted, including the large scale NTD-CTD rotation and a 5° rotation of the 8C NTD relative to the 9E NTD (**Supplementary Fig. 6A**). Notably, the AF model predicted an interaction between residues 365-382 of HROB and the MCM8C-NTD; this segment is disordered in our MCM8/9ATPγS-HROB– DNA structure, likely reflecting AF’s inability to account for the binding-induced domain rearrangements (**Supplementary Fig. 6B**). Moreover, several sidechains at the interface, including HROB N562 and MCM9 M45 and E108, adopt alternative conformations in our cryo-EM structure compared with the AF model. Interestingly and in addition to its interaction with the NTD, HROB also engages the CTD of MCM8 (*site ii*). HROB residues 398–403 insert into a surface pocket of the 8C CTD, with F398 buried in a hydrophobic cavity formed by MCM8 residues L390, I393, Y676, and L728 (**Fig. 4C & Supplementary Fig. 6C**).

To evaluate the functional importance of interfacial residues identified in the MCM8/9ATPγS-HROB–DNA structure, we constructed several HROB point mutants (F398D, M510E, H528A, F553E, and Y561E) (**Supplementary Fig. 4**) and tested their interactions with MCM8/9^680^ using *in vitro* pull-down assays (**Figure 4D-E**). MCM8/9^680^ was immobilized onto magnetic Strep-Tactin resin through an N-terminal StrepTag fused to MCM8. 6X-His-SUMO-HROB constructs are incubated with resin-bound MCM8/9^680^, followed by six washes with equal volumes of buffer. Bound complexes were subsequently eluted by addition of biotin (**Fig. 4E**).

Quantification of the eluted fractions showed that WT HROB co-eluted with MCM8 and MCM9 at relative ratios of 0.3 and 0.4, respectively, corresponding to an apparent ~1:3 stoichiometry of HROB to either MCM8 or 9, consistent with those in the cryo-EM structures. In addition to WT HROB, MCM8/9^680^ was capable of pulling down M510E and H528A, while F553E, Y561E, and F398D flowed through and were not stably retained by MCM8/9. While F553 has been previously identified and characterized by AF and biochemical assays^35^, our structures further suggest two additional bulky aromatic residues, Y561 and F398, that sit within two hydrophobic pockets at the NTD of MCM9 (*site i*) and the CTD of MCM8 (*site ii*), respectively, bridging the two tiers and potentially mediating interdomain communication within the MCM8/9 complex.

### Direct interaction between MCM8/9 and HROB is required to activate DNA unwinding

Previously, MCM8/9 was shown to bind preferentially to ssDNA and unwind fork or Holliday junction DNA substrates in a 3’ to 5’ direction, where the addition of HROB significantly stimulated both the rate and amount of DNA unwound^3,34,35^. Both His-SUMO tagged and cleaved versions of HROB^381-646^ stimulated MCM8/9 unwinding of fluorescently labeled forked DNA substrate (46 nt 3’ overhang, 10 nt 5’ flap, and 20 bp duplex with Cy3 label on DNA15 5’ end) similarly by 3-fold (**Supplementary Fig. 7A-C**).

Mutant versions of HROB^381-646^ that disrupted the interaction with MCM8/9 were examined for decreased MCM8/9 unwinding using a TBE gel-based assay to calculate apparent rates (**Fig.5A-C**). F553E displayed decreased unwinding stimulation (~2-fold) that was similar to MCM8/9 alone and consistent with a previous report^35^.

**Figure 5.**
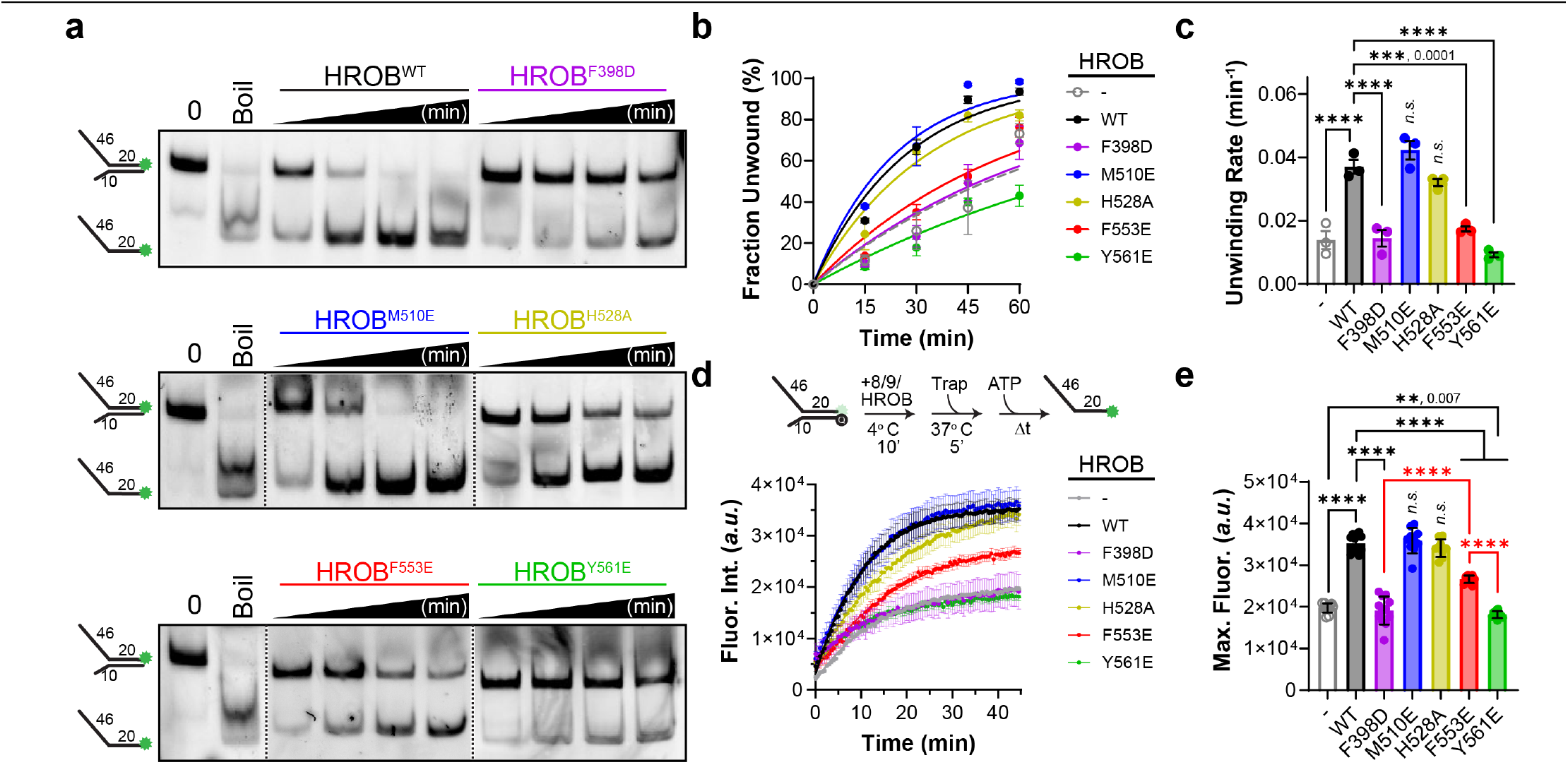
DNA unwinding activation by MCM8/9/HROB. **a**, DNA unwinding of a 3′-long arm forked DNA substrate fluorescently labeled with Cy3 resolved by TBE PAGE. The MCM8/9 hexamer is incubated with equal molar HROB constructs (colored as in **Fig. 4d-e**), initiated with ATP, and quenched at 15-minute time points. **b**, The fraction of DNA unwound at each time point was quantified, plotted, and fit to a single exponential curve. Error bars represent standard error of the mean for at least three biological replicates and are within the symbol if not visible. **c**, The unwinding rate constants, *k* (min^−1^), were plotted and compared. **d**, DNA unwinding was also measured by the increase in fluorescence (arbitrary units, *a*.*u*.) of 3′-long arm forked DNA fluorescently labeled with 5’-Cy3 opposite a 3’-black hole quencher (Q). Error bars represent standard deviation of at least eight technical replicates, and the data was fit to single exponential curve. **e**, The fitted maximum fluorescence values for each replicate were plotted and compared. Significant differences to WT HROB/MCM8/9 were determined by an ordinary one-way ANOVA where *P*-values are ****< 0.0001, explicitly provided, or determined not significant (*n*.*s*.) as indicated.

Disruption of the H-bonding residues M510E or H528A did not significantly reduce the unwinding stimulation compared to WT. However, both Y561E (at *site i*) and F398D (at *site ii*) severely disrupted DNA unwinding activation by MCM8/9 by 2.5-fold or 4-fold, respectively. In fact, it appeared that Y561E may even reduce unwinding further compared to MCM8/9 alone.

To quantify DNA unwinding in a continuous assay, similar reactions were performed to measure increasing fluorescence upon release of the 5’-Cy3 on translocating strand from the 3’ black hole quencher (Q) on the excluded strand (**Fig. 5D**). Generally, the HROB variants exhibited similar effects on MCM8/9 unwinding stimulation compared to the gel-based assays (**Fig. 5B-C** vs. **D-E**). Instead of quantifying rates of unwinding, we chose to quantify the maximum fluorescence value corresponding to the amount of unwound product. Again, F553E, Y561E, or F398D all showed significant decreases in the maximal unwound product compared the WT HROB (**Fig. 5E**). Interestingly, the maximal fluorescence for F553E was significantly higher than in the absence of HROB, and both Y561E and F398D were significantly lower than F553E, indicating that these residues (Y561 and F398) are the lynchpins connecting the MCM9 NTD to the MCM8 CTD for unwinding activation. An individual *t*-test comparing MCM8/9 on its own to addition of HROB (Y561E) shows a significant decrease with a *P*-value of 0.007.

### HROB/MCM8/9 unwinds DNA with a preference for adenine-based nucleotides

Our previous biochemical assays suggested that the γ-phosphate of ATP or ATPγS is critical for MCM8/9 loading onto DNA^3^. To understand how ATPγS binding stimulates MCM8/9 DNA binding and unwinding, we first compared the the MCM8/9^ADP^ and MCM8/9^ATPγS^ structures. Notably, the neighboring subunits within the hexameric ring move toward to each other by ~ 1.6 Å upon ATPγS binding (**Supplementary Fig. 8A and B**).

The synchronized tightening of MCM8/9 interfaces with ATPγS drives the contraction of the central channel to provide better engagement with the helical DNA, (**Supplementary Fig. 8A**), which explains the tighter binding affinity with ATPγS compared with no nucleotide or ADP alone^3^. Furthermore, we examined active site details with ADP and ATPγS. Similar as in MCM8/9^ADP^, the Walker A motif (K460 on MCM8, and K358 on MCM9) from one subunit, the sensor 2 (R711 on MCM8, and R567 on MCM9), and the arginine finger (R586 on MCM8, and R482 on MCM9) motifs from the neighboring subunit in MCM8/9^ATPγS^ contact the γ-phosphate of ATPγS (**Supplementary Fig. 9A and B**). In addition, Mg^2+^ associates between the β and γ-phosphates of ATPγS at each interfacial binding site within each of the MCM8/9^ATPγS^, MCM8/9^ATPγS^-DNA, MCM8/9^ATPγS^-HROB-DNA structures (**Supplementary Fig. 2A-C**).

Aside from the γ-phosphate group, the purine ring also plays an important role in ATP binding (**Fig. 6A-B**). The adenine base of ATPγS is observed to interact within a hydrophobic pocket (I414, V413, L605, and V609 of MCM8; M317, I501, I505, and V314 of MCM9), while the ribose group is observed to be stabilized by hydrogen bonding (Q462, E714, and T710 of MCM8; E570 of MCM9) (**Fig. 6A-B**). Previously, we have shown that any nucleoside triphosphates, including deoxynucleotides, stabilize MCM8/9 binding to ssDNA^3^. To test which (deoxy) nucleoside triphosphates support DNA unwinding by MCM8/9/HROB, we again utilized gel-based unwinding assays (**Fig. 6C**). Importantly, only the ATP within the set of ribonucleotides stimulated DNA unwinding, however interestingly, several deoxyribonucleotides also stimulated unwinding with both purines, dATP and dGTP, providing similar unwinding stimulation as ATP (**Fig. 6D**). This suggests that hydrophobic packing around the purine base, rather than specific hydrogen-bonding interactions, plays a dominant role in nucleotide selection by MCM8/9.

Furthermore, we analyzed the ATPase sites details with the higher resolution MCM8/9^ATPγS^-HROB-DNA structure (**Fig. 3B & Supplementary Fig. 9**). Within the six MCM8/9 subunits, only five ATPγS are present in MCM8/9^ATPγS^-HROB-DNA with ATPase site at the 9F/8A interface unoccupied (**Supplementary Figs. 2C & 9C**). Moreover, while the ATPase sites at 8A/9D, 8B/9E, and 8C/9F interfaces appear to be highly similar to those in MCM8/9^ATPγS^ (**Supplementary Fig. 9E, G, I**), the MCM8 subunits at the 9D/8B and 9E/8C interfaces slide around 2 Å clockwise and the 8A subunit at the ligand-free 9F/8A interface slide 2.4 Å anti-clockwise (**Supplementary Fig. 9D, F, H**). Upon this subtle ATPase domain movement, the contacting interface area between 9F and 8A reduces from 1644 Å^2^ in the MCM8/9^ATPγS^ structure to 1012 Å^2^ in the MCM8/9^ATPγS^-HROB-DNA structure. The reduced interface may enable nucleotide diffusion and subunit translocation during MCM8/9 DNA unwinding.

### The most severe mutations F398D and Y561E fail to complement MCM8/9 function in HROB knockout cells

Next, we wanted to test whether HROB mutations can complement MCM8/9/HROB function in human cells. Previously, we have shown in multiple cell lines that the addition of mitomycin C (MMC) induces nuclear foci formation of transfected GFP-MCM8 or GFP-MCM9 (**Fig. 7A**)^8,9,14,18^. Initially, we showed that knockdown of HROB with siRNA (siHROB) eliminates MMC induced nuclear foci formation of GFP-MCM9 (**Supplementary Fig. 10A)**. We then created CRISPR/Cas9 knockouts of the HROB gene in 293T cells and confirmed that GFP-MCM9 foci was eliminated with MMC treatment in two clones 3 & 5 (**Fig. 7A & Supplementary Fig.10B**). Transient transfection of a plasmid containing full-length HROB into either HROBKO clone restored MMC induced GFP-MCM9 foci as expected (**Fig. 7B & Supplementary Fig. 10C**).

Next, we transfected full-length HROB constructs with the various mutations in HROBKO cells (clones 3 or 5) to assess whether MCM9 foci formation could be restored with MMC (**Fig. 7C & Supplementary Fig. 10D**). Transfection of HROB M510E or H528A appeared to restore GFP-MCM9 foci formation similar to WT. However, neither constructs containing F398D (*site ii*), F553E (*site i*), nor Y561E (*site i*) mutations were able to restore GFP-MCM9 foci. The diffuse nuclear GFP signal appeared similar to nontreated cells or HROBKO cells with MMC. Overall, the *in vivo* complementation experiments mirror the results from the *in vitro* binding and unwinding assays (**Fig. 5**), where mutations F398D, F553E, and Y561E have the most extreme effect on MCM8/9 function.

**Figure 6.**
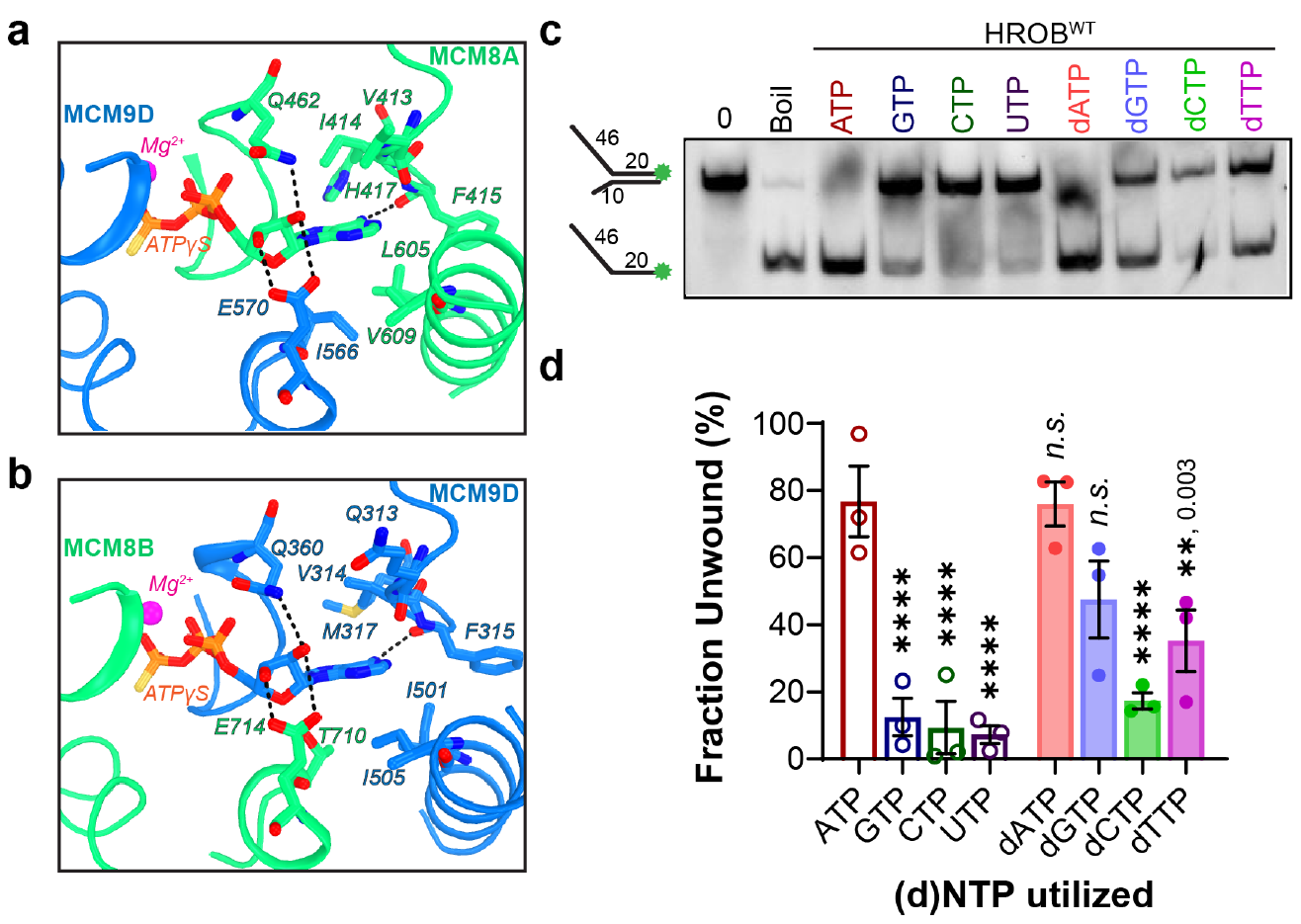
MCM8/9 prefers adenine-based (d)NTPs for DNA unwinding. Examination of the nucleotide binding pocket for **a** 9D and 8A or **b** 8B and 9D, highlighting residues interacting with ATPγS. B) DNA unwinding by HROB/MCM8/9 as in **Fig. 5a** except initiated with various (deoxy)nucleoside triphosphates and quenched at 60-minutes. **c**, The fraction unwound for three biological replicates was plotted and compared to ATP using an ordinary one-way ANOVA, where *P*-values are ****< 0.0001, explicitly provided, or determined not significant (*n*.*s*.) as indicated. Error bars represent standard error of the mean.

**Figure 7.**
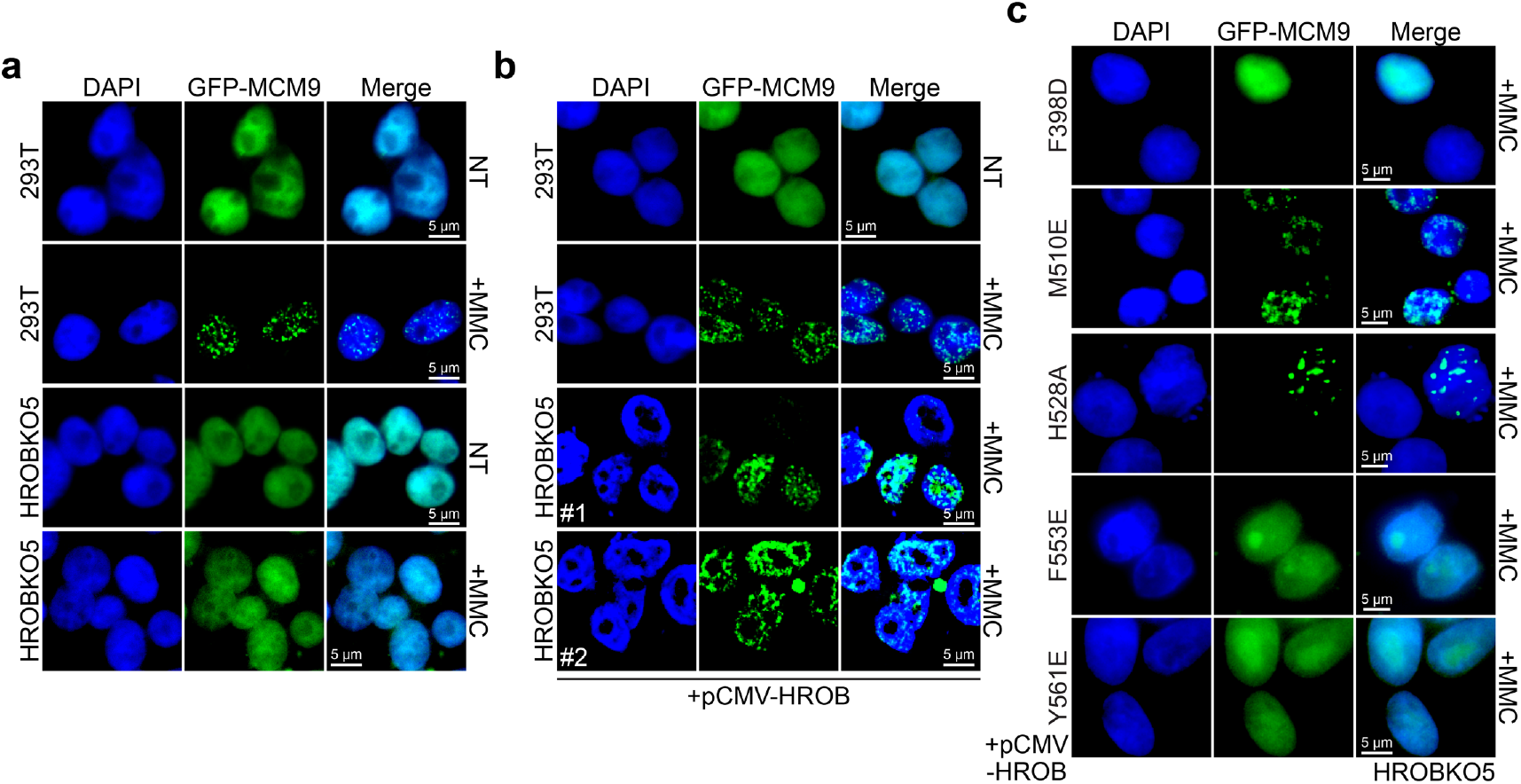
HROB interaction with MCM8/9 is required for MMC induced nuclear foci formation in human cells. **a**, GFP-MCM9 forms nuclear foci in MMC treated 293T cells but not in HROBKO cells (clone 5). **b**, Transfection of WT HROB back into HROBKO cells restores MMC induced GFP-MCM9 foci formation. #1 and #2 indicate two separate transfections. **c**, Transfection of pCMV-HROB mutants (F398D, M110E, H528A, F553E, or Y561E) into HROBKO5 to monitor any restoration of MMC induced GFP-MCM9 foci formation.

## Discussion

MCM8/9 are the most recent members of the MCM helicase family, and they are distinguished from other MCMs by a unique dimer of trimers architecture - an arrangement rarely seen among AAA^+^ helicases, aside from the recently reported heterohexameric SPATA5–SPATA5L1 peptide translocase^39^.

Our cryo-EM structures of human MCM8/9 in complex with ATPγS and DNA provide a comprehensive view of the helicase’s architecture and conformational dynamics. MCM8/9 adopts a hexameric assembly reminiscent of archaeal MCM and eukaryotic MCM2–7. Key motifs responsible for DNA binding and ATP hydrolysis are preserved across these MCM homologs. The bound DNA adopts a right-handed helical conformation and interacts with a staircase of DNA-binding loops. Notably, 9F subunit is disengaged from both ATPγS and the DNA. The reduced buried surface area at the 9F/8A interface may enable diffusion of new ATP molecule, which will drive 9F/8A association and 9F subunit translocation to the 5’-end of DNA. These observations support a rotary hand-over-hand translocation model similar to other hexameric helicases^40–44^, in which ATP hydrolysis at the 8C/9F interface triggers DNA engagement by 9F, advancing the staircase by one subunit. However, possibly because of the non-hydrolyzable ATPγS analog and/or the Walker B mutations, the ATPase sites at 8A/9D, 8B/9E, and 8C/9F appear similar as those in the MCM8/9ATPγS structure, and the specific ATPase site competent for hydrolysis relative to ssDNA binding remains unresolved.

Furthermore, we observed unique large-scale domain motions in MCM8/9 upon cofactor binding. DNA binding induces a ~28° clockwise rotation of the CTD ring relative to the NTD ring, likely reducing steric clashes and enabling subunit movement within the CTD during MCM8/9 helicase translocation and unwinding (**Fig. 8A**). A similar conformational change (26° rotation) is observed when comparing DNA-bound archaeal MCM to its apo form, although the latter is derived from a chimeric construct created from two different archaeal species (**Supplementary Fig. 3C**)^36,45^. In contrast, only a rocking motion between NTDs and CTDs has been reported for the yeast CMG complex^37,46^, and a reduced movement was observed in the human CMG^38^ (**Supplementary Fig. 3D-E**). Remarkably, binding of HROB alone is sufficient to induce a comparable NTD–CTD rotation (**Fig. 3C & 8A**). According to the structural analysis, HROB binding to the NTD of MCM9 at *site i* is likely the higher affinity site, but subsequent contact with the MCM8 CTD at *site ii* would stabilize the rotation of the CTD tier, locking it in place for DNA binding and resulting in ~3-fold stimulation in helicase activity over MCM8/9 alone.

**Figure 8.**
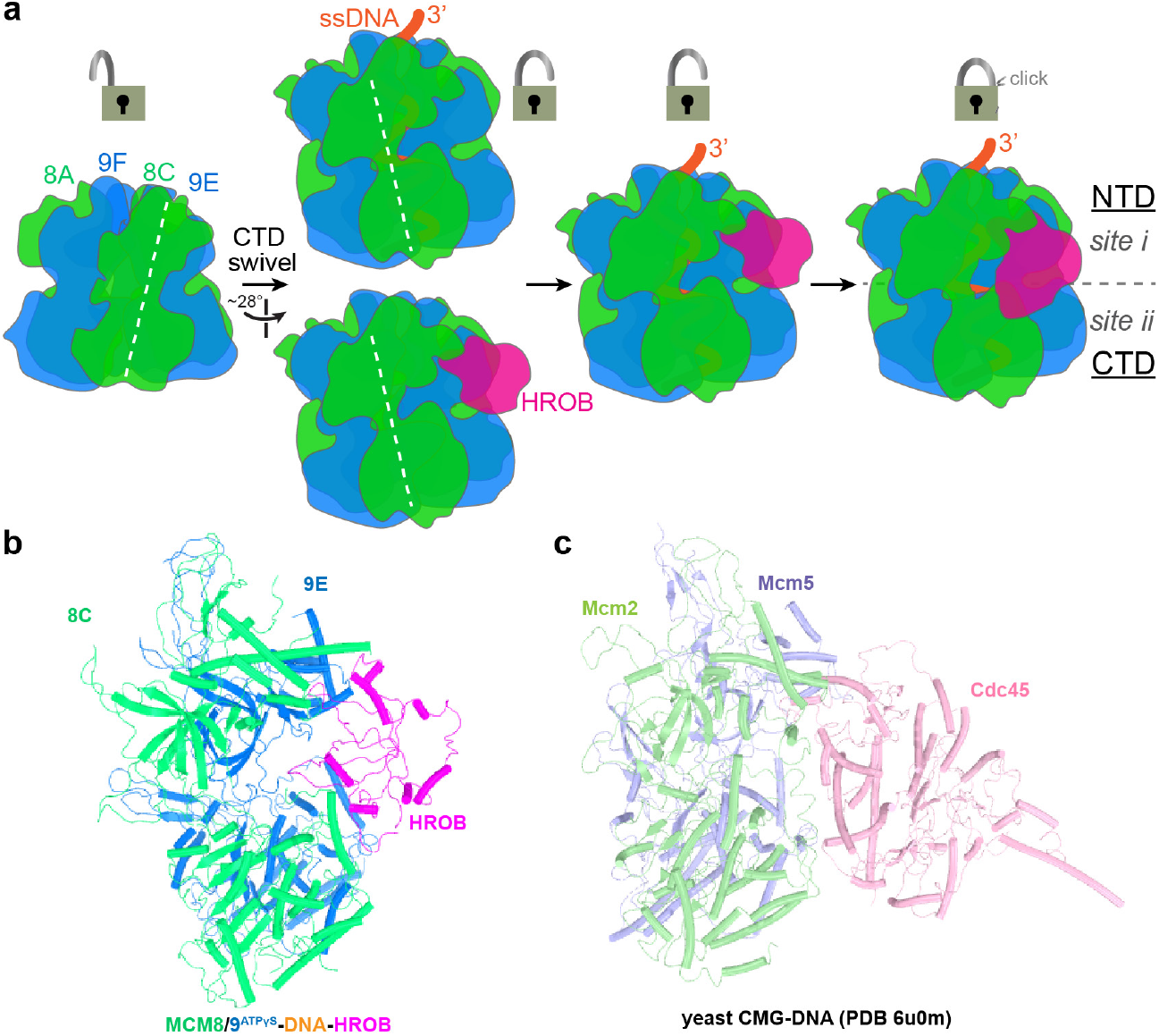
Model for MCM8/9 activation by HROB. **a**, Cartoon model of MCM8/9 DNA binding stabilization and unwinding activation by HROB. Padlocks indicate the swing of the CTD relative to the NTD, and the closed lock highlights HROB contacts at sites 1 and ii to span the NTD and CTD. Side-by-side comparison of **b** HROB binding to MCM8/9 and **c** Cdc45 binding to MCM2-7 reveals that both HROB and Cdc45 bridge the NTD and CTD to stimulate DNA unwinding.

Mutations of the bulky, aromatic residues F553 or Y561 within the OB-fold of HROB prevent binding to *site i*, while mutation of F398 prevented engagement of the CTD at *site ii* of MCM8 to lock the complex in the active state. While AlphaFold models reproduce the core HROB– MCM interfaces^35^, they failed to capture the large-scale conformational changes observed in these structures, highlighting the current limitations of predictive modeling for dynamic, multimeric assemblies.

Interestingly and still structurally unexplainable, only a single HROB binds stably to one wedge interface of MCM8/9 regardless of the availability of two other equivalent sites and having molar excesses of HROB in solution. No high quality structural classes are present with more than one HROB subunit in place. Pull-downs of HROB show ~3-fold less HROB compared to either MCM8 or MCM9. Previously, mass photometry also confirmed that HROB complexes with MCM8/9 hexamer in the same stoichiometric ratio of one MCM8/9 hexamer to one HROB monomer^35^. Additionally, it was shown that excess HROB can interfere with DNA unwinding by MCM8/9, possibly through competition of HROB and MCM8/9 for DNA binding^35^ or allosterically affecting the other two interfaces. Based on our structures, HROB stabilizes the conformation swivel of the CTD in a single state, and as the six CTDs swivel as a unit, only a single HROB is needed to connect the NTD tier to the CTD tier and lock the conformation into the active state (**Fig. 8A-B**).

Furthermore, ATPγS—but not ADP—promotes strong DNA binding^3^ resulting in a narrowing of the CTD ring around the DNA. Based on a comparison of the ADP bound^3^ and ATPγS bound structures, the γ-phosphate is important to drive the tightening and condensation of the six subunits to better engage ssDNA in the central channel for binding and translocation. However, while ATPγS is included in our cryo-EM structures, surprisingly other dNTPs can also be utilized by MCM8/9 to stabilize DNA binding and promote unwinding translocation^3^. Glutamate and glutamine side chains from adjacent MCM8 and MCM9 subunits make bifurcated hydrogen bonds to both the 2’ and 3’ OH within the AAA^+^ binding pocket not allowing directly for selection of NTPs over dNTPs. It is interesting that dATP and dGTP can stimulate unwinding similarly to that when ATP is included, suggesting that purines are favored over pyrimidines, even though GTP was unable to stimulate unwinding. Other hexameric helicases, primarily in phage and virus, have been shown to utilize multiple rNTPs and dNTPs for hydrolysis and unwinding^47–52^, and a preprint on the *E. coli* replisome goes so far to suggest that ATP is not needed at all during elongation processes^53^, however, to our knowledge, this nucleotide promiscuity has not been reported for eukaryotic hexameric helicases. Even though MCM8/9 are widely distributed in many eukaryotic supergroups, suggestive that they arose codependently early in eukaryotic evolution, they are generally absent in fungal species and some excavates, indicating that their evolutionary introduction occurred later than MCM2-7^25,54^. It is possible that nucleotide specificity had not been selected for or that possibly other MCM proteins may have similar promiscuities.

Although forked DNA substrates were utilized in the cryoEM experiments, the duplex region was not resolved in any of our structures, leaving the mechanism of strand separation and exclusion by MCM8/9 unclear. Furthermore, the NTD ring remains tightly closed in all structures, suggesting that additional factors or conformational changes may be required for MCM8/9 opening and loading onto DNA. Alternatively, MCM8/9 loading could occur through assembly of a trimer-of-heterodimers directly onto DNA. Interestingly, the overall complex of the heterodimer bound to HROB is reminiscent of Mcm2 and Mcm5 bound to Cdc45 from the CMG complex^37^, which binds the wedge interface of Mcm2 and Mcm5 and connects the NTD and CTD at Mcm2-5 (**Figure 8B-C**). Cdc45 and GINS subunits are essential auxiliary proteins and replication firing factors for converting the loaded inactive MCM2-7 double hexamer into active single CMG hexamer enzymes for bidirectional replication^55–60^. Although it is not possible to structurally align HROB and Cdc45, they act similarly in bridging the MCM tiers and activating unwinding. Further investigation is required to validate an analogous functional relationship between HROB and Cdc45 and explore the possibility that other proteins interact with MCM8/9/HROB to promote helicase loading/assembling and stimulate unwinding further similar to the GINS proteins within the CMG complex. Given the essential role of HROB for MCM8/9 activity and its synthetic lethal relationship with key cancer genes^9,33,34^, our structures may provide a foundation for rational design of therapeutics targeting the MCM8/9/HROB axis.

## Methods

### *Cloning* Hs*HROB Truncations and Mutants*

The *Hs*HROB gene (isoform 4) was synthesized (Genewiz, South Plainfield, NJ) and inserted into pET-28c plasmid with a 6X histidine-SUMO tag and a TEV cleavage site prior to the N-terminus or into pCMV3xMycTag3B with three Myc tags at the N-terminus. The pET-28c-HROB sequence was truncated to generate HROB^381-646^. HROB mutants (F398D, M510E, H528A, F553E, and Y561E) were introduced into either plasmid using the FastCloning protocol^61^ with partially overlapping primers (**Supplementary Table S1**). Plasmid sequences were screened by restriction digestion and verified using whole plasmid sequencing (Plasmidsaurus, San Francisco, CA).

### Expression and Purification of Proteins

MCM8/9^680^ expression in baculovirus infected Tni cells (Expression Systems, Davis, CA) and purification was performed as previously described^3^. The pET-28c-HROB^381-646^ plasmid constructs containing genes for truncated WT and mutant variants of *HsHROB* were transformed into Rosetta2(DE3) cells. Transformed cells were grown in LB Miller media with catabolite repressor (25% Dextrose, 25% Glycerol, 1 mM MgSO_4_, 0.1 mM MnSO_4_) overnight at 37 °C, diluted 1:50 into 2X YT media and grown to an OD_600_ between 0.6-0.8, and then induced with IPTG (1 mM) and transferred to 16 °C for 21-24 hrs. Cell pellets were harvested at 4000 RPM for 20 – 30 mins at 4 °C and stored at −80 °C. The HROB cell pellets were thawed at 4 °C and resuspended in Buffer A (50 mM Tris-HCl pH 7.5-7.6, 10% glycerol, 5 mM β-mercaptoethanol, 100 mM NaCl, 10 mM imidazole,50 mM L-arginine, 50 mM L-glutamic acid, 1 mM PMSF, 0.01% Tween 20, and 0.05 mg of benzonase). The resulting suspensions were lysed by French Press (ThermoFisher, Waltham, MA) at 1,350 PSI for 5 mins at 4 °C, repeated thrice. The lysate was clarified by centrifugation at 21k RPM at 4 °C for 1 hour and filtered through a 0.45 µm syringe filter. The clarified lysate was injected onto a pre-equilibrated 5 mL HisTrap HP column recharged with Co^2+^ attached to the AKTA Pure25 FPLC (Cytiva, Marlborough, MA) with Buffer A. The column is washed with 5 column volumes (CV) Buffer B (Buffer A with 1 M NaCl) and then an additional 5 CVs of Buffer A before eluting with an incremental stepwise gradient of Buffer A to Buffer C (Buffer A with 500 mM Imidazole). The fractions were collected and diluted with Buffer D (Buffer A with 0 mM NaCl) and then injected onto a pre-equilibrated 5 mL HiTrap Q HP column (Cytiva, Marlborough, MA) and eluted with 300 mM NaCl.

Positive fractions were combined and subjected to gel filtration using Superdex200 16/60 in buffer A to isolate a homogeneous sample which was concentrated on a pre-equilibrated 1 mL HiTrap heparin HP column (Cytiva, Marlborough, MA), eluted with 300 mM NaCl, flash frozen in liquid N_2_, and stored at −80 °C. To remove the poly-His-SUMO tag, TEV protease was added to HROB in a 1:50 ratio, incubated overnight at 4 °C, and the flow through collected from application to a 5 mL HisTrap HP Co^2+^ column. The flow through is then reconcentrated on the 1 mL HiTrap heparin column, flash frozen, and stored at −80 °C. Protein purity and size were observed through 10% SDS-PAGE analysis. The concentrations for MCM8/9, Sumo-His-HROB, and cleaved HROB were determined by A_280_ values from a Nanodrop C-1000 (ThermoFisher, Waltham, MA) using extinction coefficients 375,090, 12,950, and 9970 M^−1^cm^−1^, respectively.

### Strep-Tactin XT Pull Down Assays

Pull down assays were performed similarly as described^62^. Briefly, MCM8/9 and HROB were incubated at 4 °C at a 1:3 ratio of MCM8/9 hexamer to HROB for 10 min in wash buffer (100 mM Tris-HCl, [pH 8], 150 mM NaCl, 1 mM EDTA, 5 mM β-mercaptoethanol). Strep-Tactin XT 4Flow magnetic resin (IBA, Gottingen, Germany) is equilibrated in wash buffer with 3 washes. After incubation, proteins are added to the resin and inverted gently for 30 min at 4 °C. Resin was maintained magnetically while the flow through was collected. The resin was then washed 6 times with equal volume of wash buffer. Samples were released from the resin by incubating the resin in elution buffer (wash buffer with 50 mM D-Biotin) for 15 min at 4 °C while inverting before resolving all by SDS-PAGE staining with Simplyblue Safestain (ThermoFisher, Waltham, MA), imaged on a Gel Doc™ EZ system (Bio-Rad, Hercules, CA).The bands corresponding to MCM8, MCM9, and HROB were quantified separately using ImageQuant (Cytiva, Marlborough, MA, v.10.2), and the fractional saturation was calculated by obtaining the relative band intensity ratio of MCM9 to HROB.

### DNA Unwinding Assays

Bulk gel DNA unwinding assays were performed as previously described^3^. 5’ Cy3 labeled DNA15 was annealed to DNA232 at equimolar 10 μM concentration at 37 °C overnight in an isothermal annealing buffer (20 mM Tris–HCl [pH 8], 4% glycerol, 0.1 mM EDTA, 40 μg/ml BSA, 10 mM DTT and 10 mM MgOAc)^63^.

MCM8/9 hexamer is incubated with HROB at equal molar ratios (250 nM) with 10 nM Cy3 labelled fork DNA substrate (**Supplementary Table S1)** on ice in reaction buffer (25 mM HEPES-NaOH pH 7.5, 1 mM magnesium acetate, 1 mM dithiothreitol, 0.1mg/mL BSA) for 10 mins. The solutions were quickly warmed to 37 °C for 2.5 mins and unwinding was initiated with 5 mM ATP and 100 nM unlabeled trap (DNA15). Reactions were quenched at various timepoints (as indicated) with 5 µL quenching buffer (2% SDS, 150 mM EDTA, 30% glycerol), and then, 1 µL of 20 mg/mL Proteinase K was added. Aliquots were incubated at 37 °C for 10 min before electrophoresing on 15% TBE PAGE at 200 V for 1 hr and imaging using the RGB Typhoon (Cytiva Marlborough, MA). Quantifications of band percentage were done using ImageQuant (Cytiva Marlborough, MA) by comparing the ssDNA to dsDNA to calculate the percent unwound.

Continuous fluorescent DNA unwinding assays were performed using 250 nM MCM8/9 and HROB incubated with 50 nM of pre-annealed 5’ Cy3-DNA15 and blackhole quencher labeled DNA232-BQ (**Supplementary Table S1)** in reaction buffer for 10 min on ice. Reactions were warmed briefly to temperature of 37 °C for 5 min, and 2-fold excess of DNA15 trap and 5 mM of ATP was added prior to aliquoting technical replicates in a black 384-well microplate (Corning, Corning, NY). An increased in fluorescence corresponding to DNA unwinding was monitored over 30-sec intervals for 45 min at 37 °C using the Tecan Spark microplate reader (Tecan, Männedorf, Switzerland). The excitation wavelength was set at 510 nm and emission measured at 580 nm with monochromators set with 20 nm bandwidths and 30 flashes.

### Cryogenic Electron Microscopy Sample Preparation

The MCM8/9 heterohexamer complex was prepared as previous described^3^ with Walker B mutations (E519Q in MCM8 and E415Q in MCM9) to substantially slow down ATP hydrolysis and a C-terminal truncation of MCM9 (MCM9^680^) for cryo-EM analysis. 1 mg/mL of MCM8/9^680^ was incubated with 2-fold molar excess of hairpin fork DNA (**Supplementary Table S1)** and in the presence of 5 mM ATPγS in buffer containing 50 mM HEPES [pH 7.5], 100 mM KCl, 3 mM DTT and 5 mM MgCl_2_. After 20 min incubation at 25 °C, the reaction mixture was centrifuged at 9000 rpm for 10 min to remove any precipitate. A complex of MCM8/9^680^-HROB was prepared similarly where HROB was added in a 2-fold molar excess to the MCM8/9^680^ hexamer. 3.0 μL of freshly prepared supernatant was applied onto glow-discharged Quantifoil R1.2/1.3 Quantifoil holey carbon or UltrAuFoil holey gold R1.2/1.3 grids (Quantifoil, Großlöbichau, Thüringen, Germany). The grid was blotted for 2 s in 100% humidity at force −3 and was plunge-frozen in liquid ethane using a Vitrobot Mark IV (FEI-ThermoFisher, Hillsboro, OR).

### Data Collection and Process

For the MCM8/9/ATPγS complexes, 6,928 micrographs were collected on a Titan Krios G3 electron microscope operated at 300 kV (Laboratory for Biomolecular Structure and Dynamics (LBSD) of Texas A&M University) using the counting mode with a magnified physical pixel size of 0.832 Å with a total dose of 50 e^−^Å^−2^. The defocus values ranged between 0.6 and 2 μm. Relion 4.0 was used for initial processing of the data.

MotionCor2 was used for drift-correction and electron-dose-weighting for all movies^64^. The defocus values were estimated on non-dose-weighted micrographs with Gctf^65^. 3,184,098 particles were picked with template-free picking in Relion and then imported in cryoSPARC^66^ for subsequent 2D classification, 3D classification, and heterogenous refinement. The classes showing clear secondary structures’ features and with or without DNA were separately processed with additional 3D classification and heterogenous refinement in cryoSPARC. Following non-uniform refinement in cryoSPARC, the particles were subjected to Ctf-refinement and polishing in Relion. The final maps are generated with the polished particles using non-uniform refinement in cryoSPARC to 3.51 Å (MCM8/9^680^/ATPγS) and 3.58 Å (MCM8/9^680^/ATPγS/DNA) by the 0.143 cutoff criterion in FSC.

For the MCM8/9^680^-HROB complexes, 15,244 micrographs were collected on a Titan Krios GIF electron microscope operated at 300 kV (NYSBC) using the counting mode with a nominal magnification of 165 K (calibrated pixel size of 0.832 Å). Similarly, motion-correction and ctf estimation were performed with MotionCor2 and Gctf in Relion. 7,765,979 particles were picked with template-free picking. Following two rounds of 2D classification and 3D classification in cryoSPARC^66^, the classes showing clear secondary structures’ features and with or without DNA were separately processed with additional 3D classification and heterogenous refinement in cryoSPARC^66^. The final map of MCM8/9/ATPγS/HROB complex without DNA was generated at 4.16 Å with non-uniform refinement in cryoSPARC. For the MCM8/9/ATPγS/DNA/HROB complex, the particles were subjected to Ctf-refinement and Polishing in Relion and the final maps are generated with the polished particles using non-uniform refinement in cryoSPARC to 2.96 Å. To better refine the HROB portion of the complex, a local mask was generated with CHIMERA and used for particle signal subtraction and local 3D classification in cryoSPARC. The class showing the most complete HROB was refined with non-uniform refinement in cryoSPARC to 3.72 Å.

### Model Building and Refinement

We used our previous MCM8/9 structure^3^ as initial models for model building. The structure of DNA was generated with COOT and the structure of HROB was generated with AlphaFold. Each of the protein chains were manually docked into the cryo-EM density maps in Chimera^67^. The models were first manually adjusted in COOT^68^ and then refined in Phenix^69^ with real-space refinement and secondary structure and geometry restraints. For MCM8/9/ATPγS/HROB complex without DNA, only rigid-body refinement was performed in Phenix considering the poor local resolution of the CTD domains. Statistics of all cryo-EM data collection and structure refinement are shown in **Supplementary Table S2**.

### Generation of HROB Knockout Cells

293T cells were cultured in Dulbecco’s Modified Eagle Medium (DMEM) (Gibco, Waltham, MA) supplemented with 10% fetal bovine serum (FBS) (Gibco) at 37 °C in a humidified incubator with 5% CO_2_. Knockout of the HROB gene (HROB^KO^) was performed in parental 293T cells using CRISPR-Cas9 gene disruption. A guide RNA (gRNA) sequence (5’-GGACCTTTTGGGGTACCAAC) targeting exon 3 of the HROB gene was selected and cloned into the pSpCas9(BB)-2A-Puro (PX459) V2.0 vector (Addgene #62988) using Platinum^™^ SuperFi^™^ II DNA Polymerase (ThermoFisher, Waltham, MA) and associated partially overlapping primers (**Supplementary Table S1**). 293T cells were transfected with pSpCas9(BB)-2A-Puro-gHROB using LPEI (Fisher Scientific, Waltham, MA) in reduced serum Opti-MEM media (Gibco, Waltham, MA). After 48 hours, cells were transferred to selection media containing 1 μg/ml puromycin (ThermoFisher, Waltham, MA) and cultured for an additional 72 hours with fresh puromycin containing media replaced daily to maintain selection and promote enrichment of transfected populations. Single cells were sorted into 96-well plates using an array dilution method. Established clones were phenotypically screened for a reduction in MMC induced pEGFP-MCM9 nuclear foci.

### Cell Transfections

Parental 293T or HROB^KO^ cells were seeded onto glass coverslips coated with poly-L-lysine (Newcomer Supply, Waunakee, WI) and co-transfected with pEGFP-MCM9 and each 3X-Myc tagged HROB mutant plasmids using LPEI. Briefly, plasmid DNA and LPEI were mixed (1:3 ratio) and incubated at room temperature for 20 minutes before being added to cells in reduced serum Opti-MEM media. For immunofluorescence experiments, six hours after transfection, the medium was replaced with complete DMEM/10% FCS media, and cells were incubated overnight. The following day, cells were treated with 3 μM mitomycin C (MMC) (Cell Signaling Technology, Danvers, MA) for 6 hours and then processed for immunofluorescence.

### Immunofluorescence Microscopy

Cells grown on coverslips were pre-extracted with 0.5% Triton X-100 for 1 min and fixed in 4% paraformaldehyde for 15 minutes at room temperature, permeabilized with 0.1% Triton X-100 in 1X PBS for 10 minutes. Coverslips were then washed three times with 1X PBS. Cells were mounted in mounting media containing DAPI (ProLong Diamond Antifade Mountant with DAPI, ThermoFisher, Waltham, MA) and sealed with clear nail polish and imaged under a FV-1000 epifluorescence laser scanning microscope (Olympus Corp., Center Valley, PA). Images were processed with Fluoview (Olympus, v.4.2b) or cellSens software (Olympus, v2).

## Data availability

Cryo-EM maps and atomic models were deposited to the EM Data Bank and PDB under accession numbers EMD-XXXXX and YYY for MCM8/^9ATPγS^, EMD-XXXXX and YYY for MCM8/^9ATPγS^/DNA, EMD-XXXXX and YYY for MCM8/^9ATPγS^/HROB, EMD-XXXXX and YYY for MCM8/^9ATPγS^/DNA-HROB, and EMD-XXXXX and YYY for locally improved MCM8/^9ATPγS^/DNA-HROB respectively. Data and materials can be obtained from the corresponding authors upon request. Source data are provided with this paper.

## Supplementary Data

Supplemental material: Tables S1 and S2 and Figures S1-S10 are available at ??? online.

## Acknowledgements

We thank all members of the Gao and Trakselis laboratories for productive conversations and insights. We acknowledge the Baylor Molecular Bioscience Center (MBC) for providing instrumentation and resources aiding this project. We thank Dr. Gaya P. Yadav at the Laboratory for Biomolecular Structure and Dynamics (LBSD) of Texas A&M University and National Center for Cryo-EM Access and Training (NCCAT) for cryo-EM sample screening and data collection. The LBSD is supported, in part, by the Department of Biochemistry & Biophysics, AgriLife, and Texas A&M University.

## Author contributions

C.L performed the cryo-EM analysis and structural modelling. C.T cloned all of the HROB constructs, performed the protein purifications and *in vitro* binding and unwinding assays. T.M.A created the HROB knockout cells and performed all of the cell transfections and fluorescence microscopy. D.R.M. cloned, expressed and purified the original MCM8/9 constructs. C.L., C.T., T.M.A, D.R.M, Y.G. and M.A.T. reviewed and participated in the data analysis and interpretation as well as writing, editing, and reviewing the manuscript.Y.G. and M.A.T. supervised the project and provided funding support.

## Abbreviations

AAA^+^: ATPases Associated with diverse cellular Activities
AF: AlphaFold
bp: base pair
CMG: Cdc45-GINS-MCM2-7
cryo-EM: cryogenic electron microscopy
CTD: C-terminal domain
CTE: C-terminal extension
CV: column volume
dsDNA: double stranded
DNA; h2i: helix 2 insertion
HR: homologous recombination
HROB: homologous recombination OB-fold
ICL: interstrand crosslink
MCM: Minichromosomal maintenance
MMC: mitomycin C
MMR: mismatch repair
NTD: N-terminal domain
NTPs: nucleotide triphosphates
POF: primary ovarian failure
POI: primary ovarian insufficiency
ps1β: presensor 1 beta
RMSD: root-mean-square deviations
ssDNA: single-stranded DNA
*Sso*MCM: *Saccharolobus solfataricus* MCM
ZF: zinc finger

