## Supplementary for "Structural Activation of DNA Unwinding by MCM8/9/HROB"

#### Author ORCIDs

Chuxuan Li ORCID: 0000-0001-9386-9909
Colin To ORCID: 0009-0003-4462-1423
Temitope M. Adeleke ORCID: 0009-0008-7041-6921
David R. McKinzey ORCID: 0000-0003-0845-5352
Yang Gao ORCID: 0000-0002-4037-0431
Michael A. Trakselis ORCID: 0000-0001-7054-8475

### Supplementary Tables

#### Supplementary Table S1: DNA plasmids, primers, and substrates

| Plasmids | Source |
|---|---|
| pUC57_Kan-HROB | Genewiz |
| pFastBacDual-MCM8/9$^{680}$ | McKinzey *et al.* 2023 |
| pET28c | Clontech |
| pET28c-SUMO-TEV-HROB (Iso4) | This work |
| pET28c-SUMO-TEV-HROB$^{381-646}$ | This work |
| pET28c-SUMO-TEV-HROB$^{381-646}$(F398D) | This work |
| pET28c-SUMO-TEV-HROB$^{381-646}$(M510E) | This work |
| pET28c-SUMO-TEV-HROB$^{381-646}$(H528A) | This work |
| pET28c-SUMO-TEV-HROB$^{381-646}$(F553E) | This work |
| pET28c-SUMO-TEV-HROB$^{381-646}$(Y561E) | This work |
| pCMV-Tag3B | Clontech |
| pCMV-3xMycTag3B | This work |
| pCMV-3xMycTag3B-HROB | This work |
| pCMV-3xMycTag3B-HROB(F398D) | This work |
| pCMV-3xMycTag3B-HROB(M510E) | This work |
| pCMV-3xMycTag3B-HROB(H528A) | This work |
| pCMV-3xMycTag3B-HROB(F553E) | This work |
| pCMV-3xMycTag3B-HROB(Y561E) | This work |
| pEGFP-MCM9 | McKinzey *et al.* 2023 |
| pSpCas9(BB)-2A-Puro(PX459) | Addgene #62988 |
| pSpCas9(BB)-2A-Puro-gHROB | This work |

| Primers | Sequence (5'-3') |
|---|---|
| HROBΔQ484F | GAAATAATTTTGTTTAACTTTAATAAGGAGATATACCATGGCGTGCAGTTTGCAG |
| HROBΔQ484R | GGCAGCAGCCTAGGTTAATTAATTTTTCAAATTGGGGGTGTGACCAACTACTGGTCCC |
| HROB381ForBsiWI | CTTTATTTTCAGGGCCGTACGCCCCAGCAGCCCACCCATC |
| pET28cHisTEVHROB_Rev | GCCCTGAAAATAAAGATTCTCGAATTG |
| HROB(F398D)For_SmaI | CCGTGACCCGGGCCCAGCTGGGATCCTGCCTCACCAGC |
| HROB(F398D)Rev_SmaI | CTGGGCCCGGGTCACGGCGAGTTTTGGCTCGGGTGGAGGG |
| HROB(M510E)For_ScaI | CGGAGTACTGAAGACGCCAGTGTGGTTTTCAAGGACC |
| HROB(M510E)Rev_ScaI | GCGTCTTCAGTACTCCGAGTCAGGGACTTGATCATCAC |
| HROB(H528A)For_MscI | GGACGGTGGCCAGGTTGCTGCTGGAGACGTGCCAGAATGAG |
| HROB(H526A)Rev_MscI | CAACCTGGCCACCGTCCCCTGCATCTCTCCCGTGGGG |
| HROB(F553E)For BstB1 | GGAGTCTCCACTCCGAAATCACTACCTCAACGTGAC |
| HROB(F553E)Rev BstB1 | GGAGTGAAGGAGACTCCCACTCCAATCTGCTTCAGCAGCAG |
| pCMV3XTagBFor | AGCCCGGGCGGATCCCCCG |
| pCMV3XTag3BRev | CTTCTAAATCTCGTTTTCAATTAAAGGCTCCTCCTGAAT |
| HROBpCMVFor | TTTCCGAGGAGGACTTAATGGCGTGCAGTTTGCAGAAGCTGTTTG |
| HROBpCMVRev | GGGGATCCGCCCGGGCTCTAACTACTGGTCCCACAGAAGAAGTCTTC |
| pCMVTag3BFor-3X | CAGTGAAGAAGATTTAGAGCAAAAGTTAATTTCCGAGGAGGACTTAAGCCCGGGCGGATCCCCCG |
| pCMVTag3BRev-3X | TAAATCTTCTTCACTGATTAGTTTCTGTTCCAGATCCTCTTCAGAGATGAGTTTCTGCTC |
| pCMVTag3XBFor | AGCCCGGGCGGATCCCCCG |
| pCMVTag3XBRev | CTTCTAAATCTCGTTTTCAATTAAAGGCTCCTCCTGAAT |
| HROBpCMVFor | TTTCCGAGGAGGACTTAATGGCGTGCAGTTTGCAGAAGCTGTTTG |
| HROBpCMVRev | GGGGATCCGCCCGGGCTCTAACTACTGGTCCCACAGAAGAAGTCTTC |
| gHROBFor | CCTTTTGGGGTACCAACGTTTTAGAGCTAGAAATAGCAAGTTAAAATAAGGCTAG |
| gHROBRev | TGGTACCCCAAAAGGTCCGGTGTTTCGTCCTTTCCACAAGATATATAAAGC |

| DNA Substrates | Sequence (5'-3') |
|---|---|
| Hairpin Fork | TTTTTTTTTTCGTTCCGGTTTCCGGAACGTTTTTTTTTTTTTTTTTTTTTTTT |
| DNA15Cy3 | Cy3-CACCTCTCCCTACGCTTCCCACCCACCCCGACCGGCATCTGCTATGGTACGCTGAGCGAGAGTAGC |
| DNA15 | CACCTCTCCCTACGCTTCCCACCCACCCCGACCGGCATCTGCTATGGTACGCTGAGCGAGAGTAGC |
| DNA232_BHQ | TTTTTTTTTTGGGAAGCGTAGGGAGAGGTG-BHQ |
| DNA232 | TTTTTTTTTTGGGAAGCGTAGGGAGAGGTG |

| siRNA | Source |
|---|---|
| HROB | Sigma - SASI_Hs02_00356457; SASI_Hs01_00035908 |

**Table S2. Cryo-EM data collection, refinement, and validation statistics**

| | MCM8/9$^{ATP\gamma S}$ | MCM8/9$^{ATP\gamma S}$-DNA | MCM8/9$^{ATP\gamma S}$-DNA-HROB | MCM8/9$^{ATP\gamma S}$-HROB | MCM8/9$^{ATP\gamma S}$-DNA-HROB (local) |
|---|---|---|---|---|---|
| **Data collection and processing** | | | | | |
| EMDB ID | | | | | |
| PDB ID | | | | | |
| Microscope | Krios | Krios | Krios | Krios | Krios |
| Detector | Gatan K3 | Gatan K3 | Gatan K3 | Gatan K3 | Gatan K3 |
| Magnification | 165000 | 165000 | 165000 | 165000 | 165000 |
| Voltage (kV) | 300 | 300 | 300 | 300 | 300 |
| Electron exposure ($e^-$/ $Å^2$) | 49 | 49 | 50 | 50 | 50 |
| Defocus range (μM) | 0.6-3 | 0.6-3 | 0.6-2 | 0.6-2 | 0.6-2 |
| Pixel size (Å) | 0.832 | 0.832 | 0.83 | 0.83 | 0.83 |
| Number of movies | 6928 | | | 15244 | |
| Initial particles (no.) | 3184098 | | | 7765979 | |
| Final particles (no.) | 123685 | 53401 | 97305 | 23773 | 7223 |
| Map resolution (Å) | 3.51 | 3.58 | 2.96 | 4.16 | 3.72 |
| FSC threshold | 0.143 | 0.143 | 0.143 | 0.143 | 0.143 |
| **Refinement and Validation** | | | | | |
| No. of atoms | 28895 | 29765 | 30742 | 30140 | 31056 |
| No. of residues | | | | | |
| Protein | 3651 | 3750 | 3870 | 3820 | 3913 |
| Nucleic acid | 0 | 10 | 13 | 0 | 12 |
| Ligand | 12 | 10 | 10 | 10 | 10 |
| R.M.S. deviations | | | | | |
| Bond lengths (Å) | 0.002 | 0.002 | 0.002 | 0.002 | 0.002 |
| Bond angles (°) | 0.570 | 0.547 | 0.516 | 0.656 | 0.535 |
| Clashscore | 6.41 | 6.76 | 8.26 | 8.86 | 8.79 |
| MolProbity | 2.18 | 2.28 | 2.38 | 2.27 | 2.43 |
| Correlation coefficients | | | | | |
| CC (mask) | 0.70 | 0.80 | 0.78 | 0.72 | 0.78 |
| CC (peaks) | 0.66 | 0.73 | 0.70 | 0.65 | 0.62 |
| CC (volume) | 0.71 | 0.78 | 0.77 | 0.71 | 0.76 |
| Ramachandran plot (%) | | | | | |
| Favored | 92.16 | 91.76 | 91.85 | 92.42 | 91.71 |
| Allowed | 7.59 | 8.02 | 7.97 | 7.29 | 8.11 |
| Outliers | 0.25 | 0.22 | 0.18 | 0.29 | 0.18 |

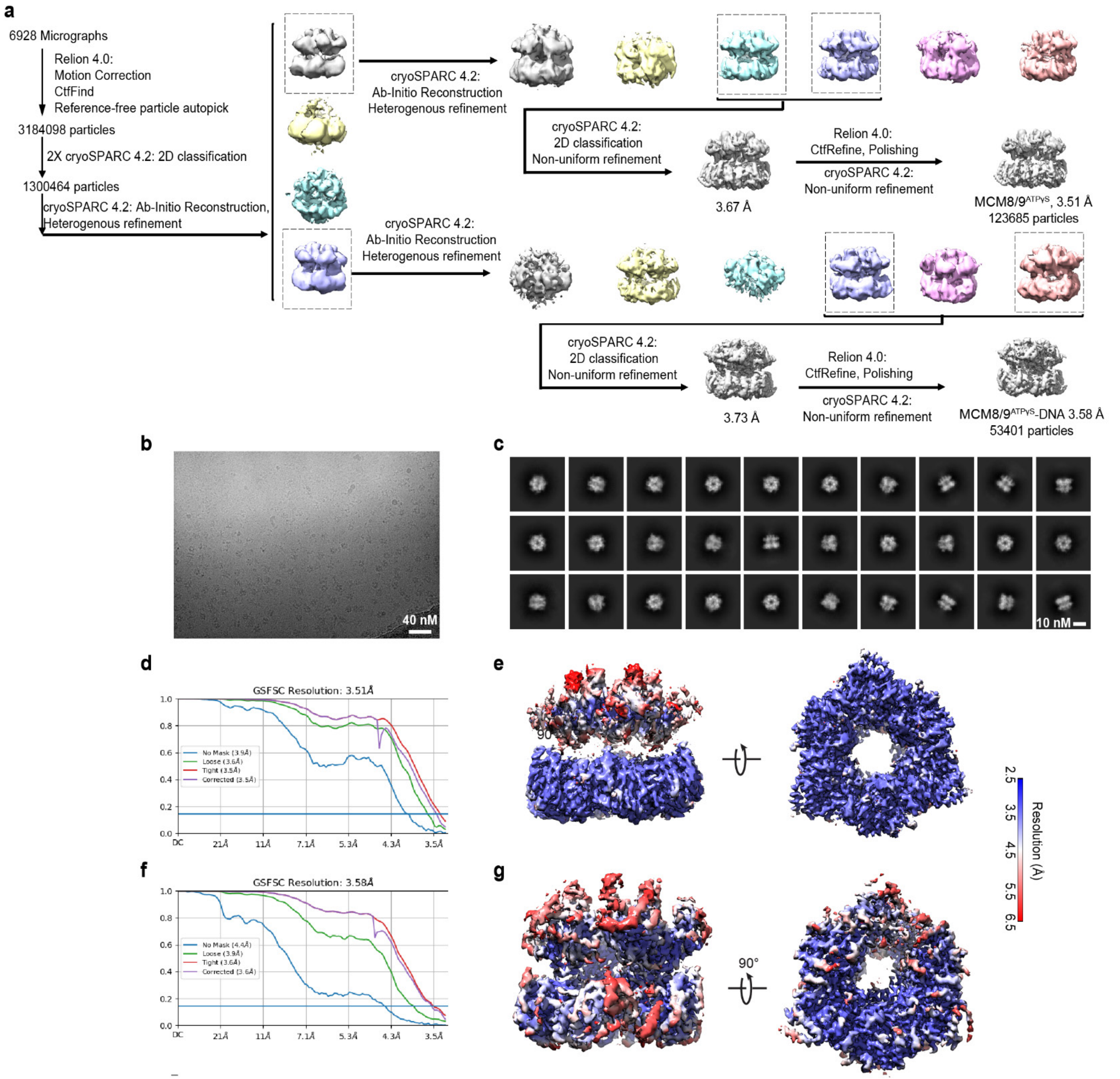


**Supplementary Fig. 1. Cryo-EM data processing of MCM8/9-DNA complex.**
**a,** Overview of image processing and refinement strategy. **b,** A representative cryo-EM micrograph. **c,** 2D class averages of MCM8/9-DNA complex. **d,** FSC curves between the two half maps with indicated resolution at FSC = 0.143 for the MCM8/9$^{ATP\gamma S}$ complex. **e,** Local resolution estimation of the cryo-EM density map for the MCM8/9$^{ATP\gamma S}$ complex. **f,** FSC curves between the two half maps with indicated resolution at FSC = 0.143 for the MCM8/9$^{ATP\gamma S}$-DNA complex. **g,** Local resolution estimation of the cryo-EM density map for the MCM8/9$^{ATP\gamma S}$-DNA complex.

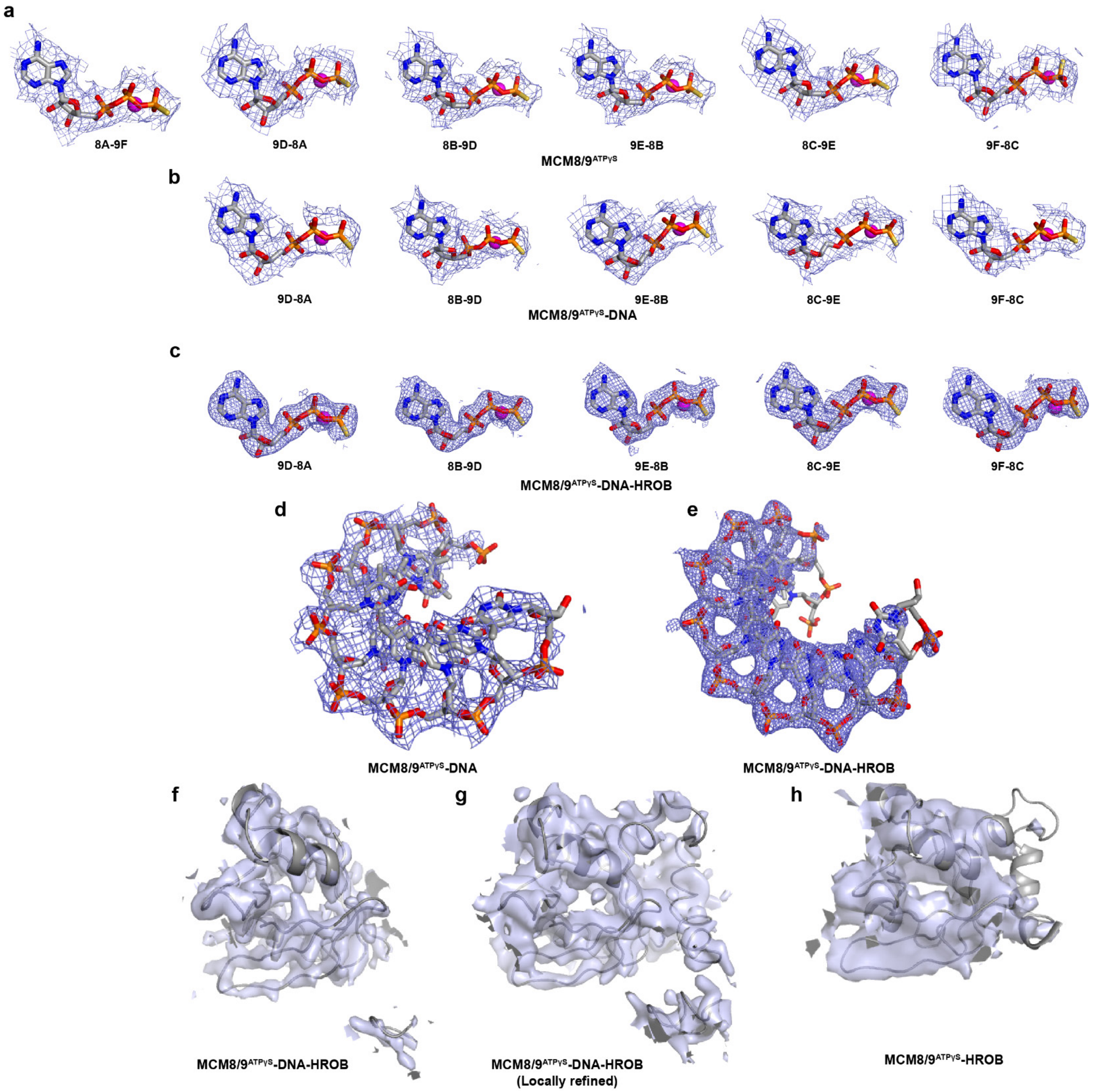


**Supplementary Fig. 2. Local cryo-EM maps of the MCM8/9 complexes.**
Local cryo-EM maps of ATPγS and $Mg^{2+}$ in MCM8/9$^{ATPγS}$ **a,** MCM8/9$^{ATPγS}$ DNA **b,** and MCM8/9$^{ATPγS}$-HROB-DNA **c,** structures. The ATPγS, $Mg^{2+}$, and their associated maps were shown as stick, sphere, and mesh, respectively. Local cryo-EM maps of DNA in **d,** MCM8/9$^{ATPγS}$ DNA and **e,** MCM8/9$^{ATPγS}$-HROB-DNA structures. The DNA and their associated maps were shown as stick and mesh, respectively. Local cryo-EM maps of **f,** HROB in MCM8/9$^{ATPγS}$-HROB-DNA, **g,** locally refined MCM8/9$^{ATPγS}$-HROB-DNA, **h,** MCM8/9$^{ATPγS}$-HROB structures. The HROB subunits and their associated maps were shown as cartoon and transparent surface, respectively.

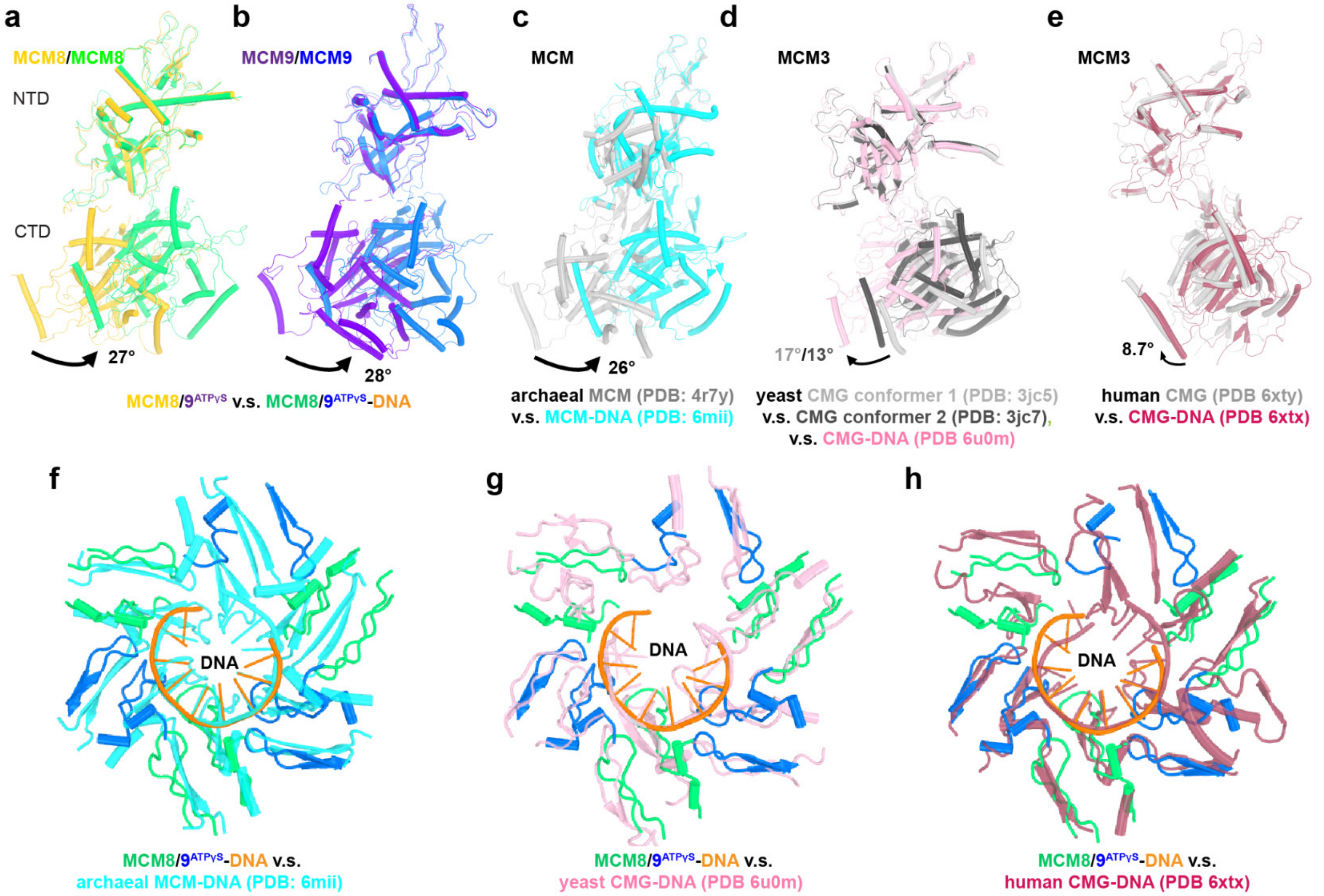


**Supplementary Fig. 3. DNA binding and domain rotations in MCM8/9 and its homologs.**
Domain movement upon DNA binding in MCM8 **a,** MCM9 **b,** archaeal MCM **c,** yeast MCM3 subunit **d,** and human MCM3 subunit **e**. In all panels, the NTD of MCM subunits were aligned and the CTD movements were indicated by black arrows. The **a** and **b** panels are color-coded as in **Figure 1**. In **c-e**, the DNA binding structures were colored in cyan, pink, and salmon for archaeal, yeast, and human MCM subunits, respectively. The archaeal MCM without DNA was derived from a chimeric MCM with NTD from *P. furiosus* MCM and CTD from *S. solfataricus* MCM, whereas the *S. solfataricus* MCM was used for the archaeal MCM-DNA complex. **d,** The yeast CMG structures were captured in two conformations, shown in light and dark grey. Structural alignments of MCM8/9 DNA binding loops with those in **f,** archaeal MCM, **g,** yeast CMG, and **h,** human CMG.

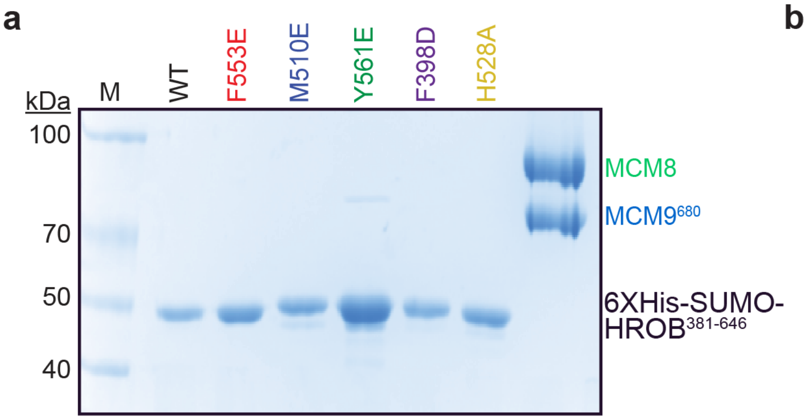


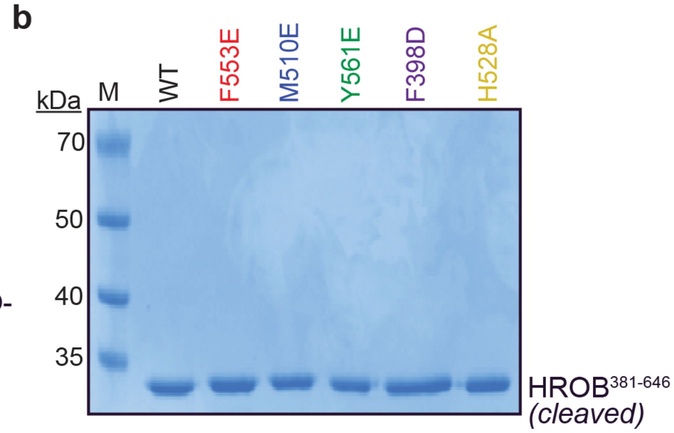


**Supplementary Fig. 4. Purification of proteins.**
**a,** An SDS-PAGE gel of all 6X-SUMO tagged HROB variants and MCM8/9$^{680}$. **b,** The tags were cleaved with TEV protease providing the cleaved versions of HROB.

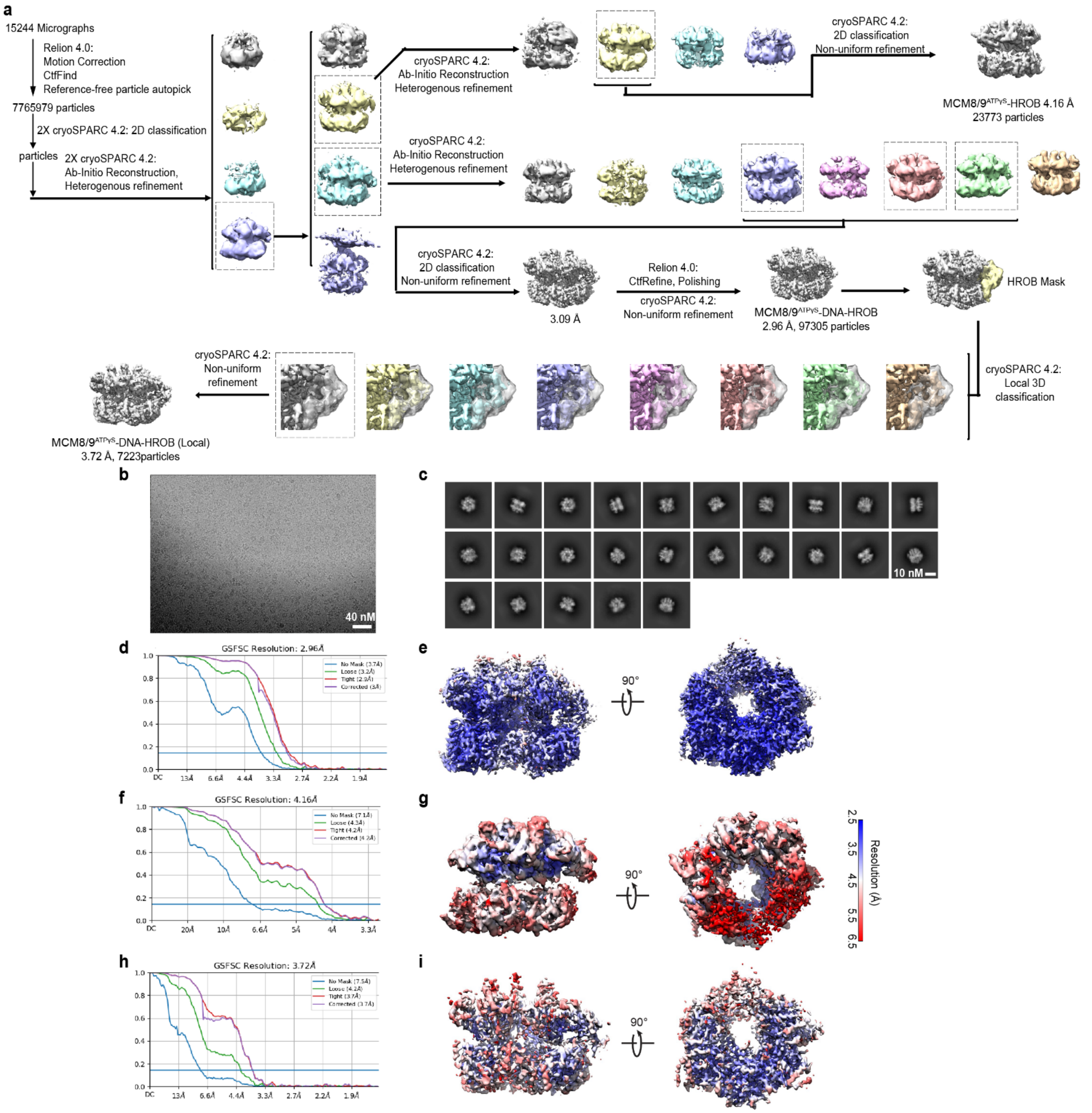


**Supplementary Fig. 5. Cryo-EM data processing of MCM8/9-HROB-DNA complex.**
**a,** Overview of image processing and refinement strategy. **b,** A representative cryo-EM micrograph. **c,** 2D class averages of MCM8/9-HROB-DNA complex. **d,** FSC curves between the two half maps with indicated resolution at FSC = 0.143 for the MCM8/9$^{ATP\gamma S}$-HROB-DNA complex. **e,** Local resolution estimation of the cryo-EM density map for the MCM8/9$^{ATP\gamma S}$-HROB-DNA complex. **f,** FSC curves between the two half maps with indicated resolution at FSC = 0.143 for the MCM8/9$^{ATP\gamma S}$-HROB complex. **g,** Local resolution estimation of the cryo-EM density map for the MCM8/9$^{ATP\gamma S}$-HROB complex. **h,** FSC curves between the two half maps with indicated resolution at FSC = 0.143 for the locally refined MCM8/9$^{ATP\gamma S}$-HROB-DNA complex. **i,** Local resolution estimation of the cryo-EM density map for the locally refined MCM8/9$^{ATP\gamma S}$-HROB-DNA complex.

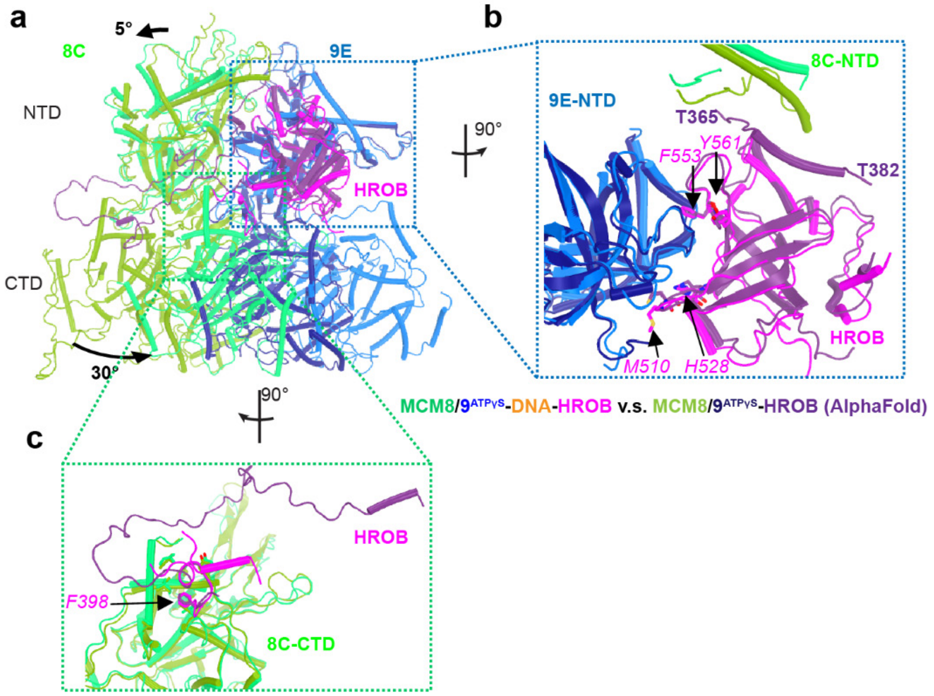


**Supplementary Fig.6. MCM8/9 interaction with HROB compared with AlphaFold model.**
**a,** Comparison of MCM8/9 interaction with HROB in our MCM8/9$^{ATP\gamma S}$-HROB-DNA structure and AlphaFold predicted model. The NTD of 9E subunit was aligned and the domain rotations of 8C-NTD and CTDs were indicated by black arrows. Zoomed in view of HROB interaction with **b,** MCM8/9 NTD and **c,** CTD.

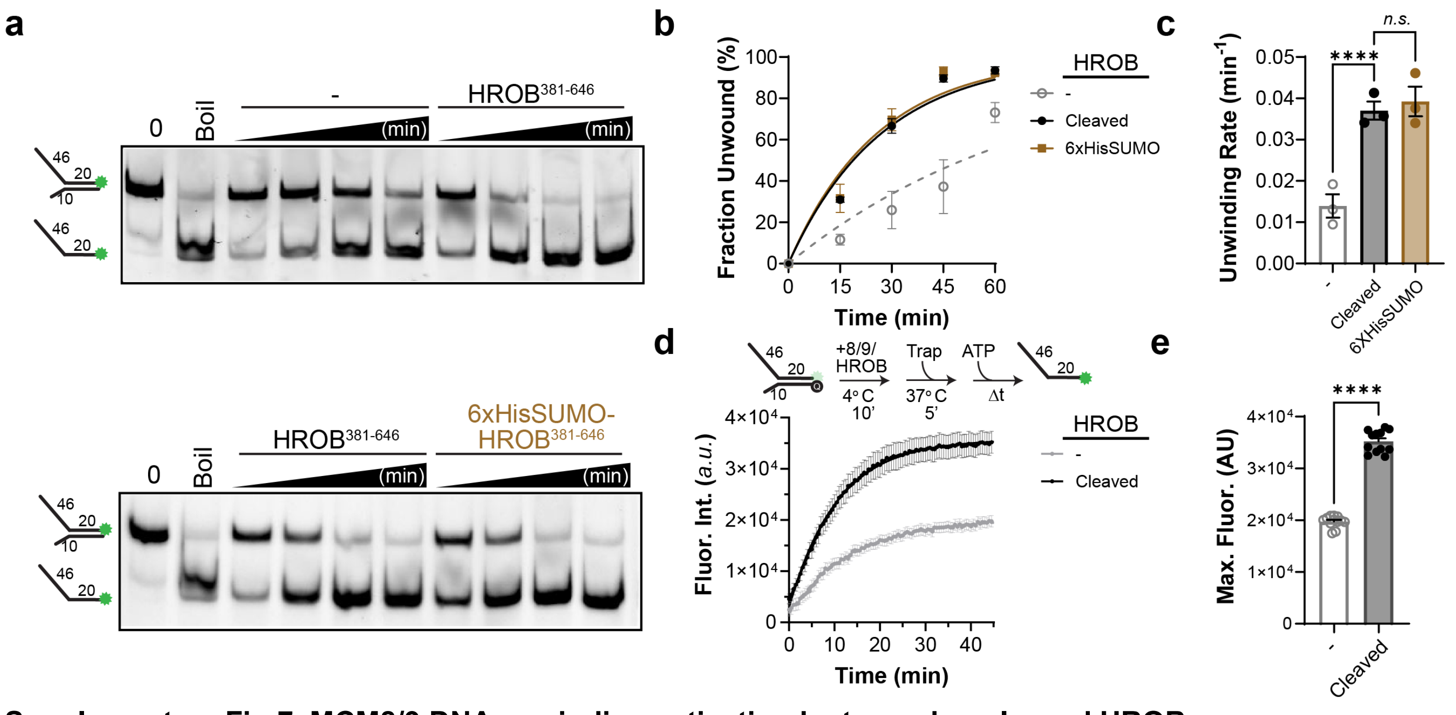


**Supplementary Fig.7. MCM8/9 DNA unwinding activation by tagged or cleaved HROB.**
**a,** TBE PAGE gels of DNA unwinding by HROB/MCM8/9 as in **Fig. 5A** with either TEV cleaved HROB$^{381-646}$ or uncleaved 6xHisSUMO-HROB$^{381-646}$ (**Extended Data Fig. 2**). **b,** The fraction of DNA unwound at each time point was quantified, plotted, and fit to a single exponential curve. Error bars represent standard error of the mean for at least three biological replicates and are within the symbol if not visible. **c,** The unwinding rate constants, *k* ($min^{-1}$), were plotted and compared. **d,** DNA unwinding was also measured by the increase in fluorescence (arbitrary units, *a.u.*) of 3′-long arm forked DNA fluorescently labeled with 5’-Cy3 opposite a 3’-black hole quencher (Q). Error bars represent standard deviation of at least eight technical replicates, and the data was fit to single exponential curve. **e,** The fitted maximum fluorescence values for each replicate were plotted and compared. Significant differences to WT HROB/MCM8/9 were determined by an ordinary one-way ANOVA (*n.s.* = not significant, ****$P < 0.0001$).

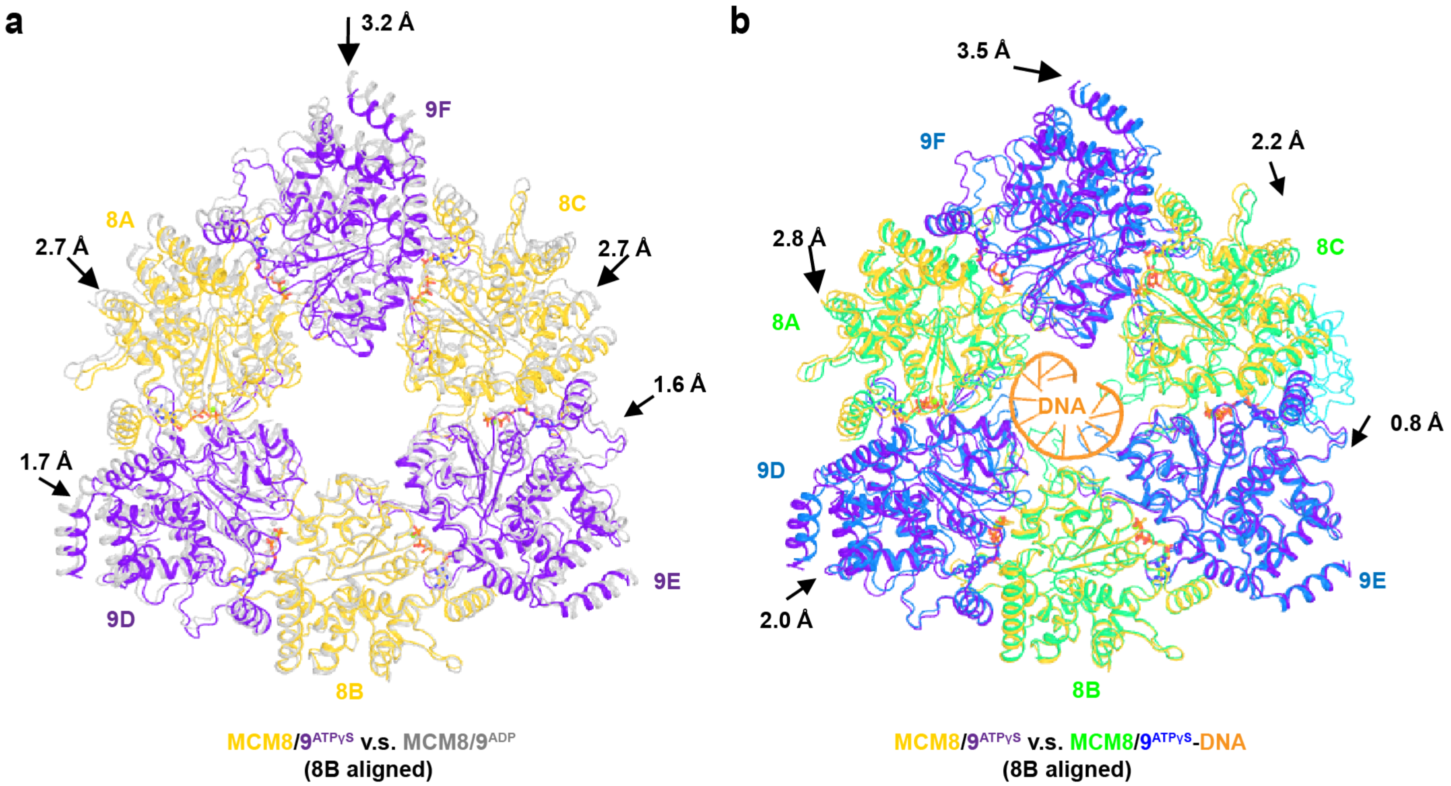


**Supplementary Fig. 8. ATP and DNA binding induced domain movement in the MCM8/9 CTDs.**
**a,** Comparison of CTDs between MCM8/9$^{ATPγS}$ (gold and purple) and MCM8/9$^{ADP}$ (grey, PDB ID 8S91). The 8B subunit was aligned and the movements of other domains relative to 8B were indicated by black arrows. **b,** Comparison of CTDs between MCM8/9$^{ATPγS}$ (gold and purple) and MCM8/9$^{ATPγS}$-HROB-DNA (green and blue). The 8B subunit was aligned and the movements of other domains relative to 8B were indicated by black arrows.

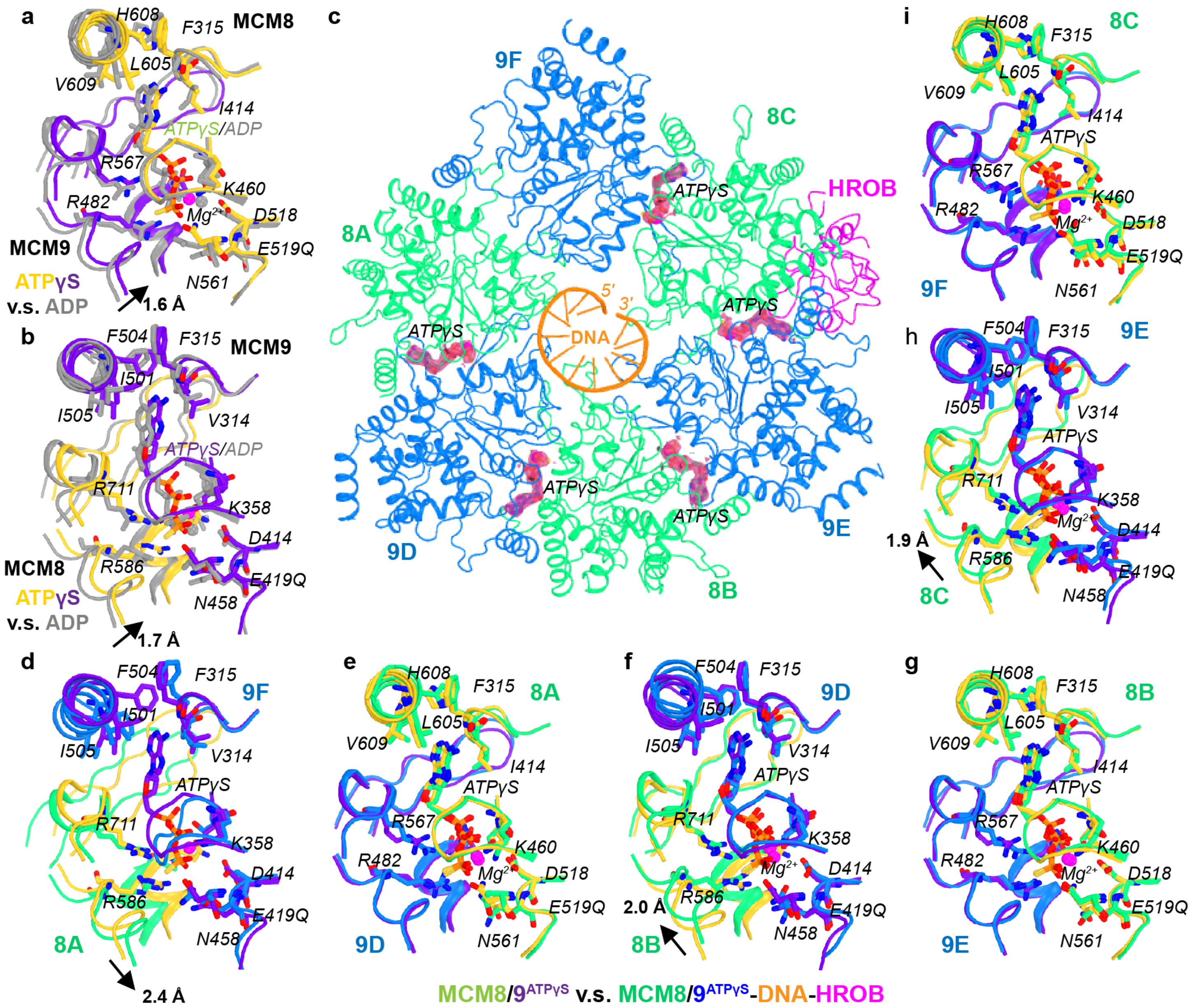


**Supplementary Fig. 9. ATPase site configurations in MCM8/9 complexes.**

**a-b,** Comparison of ATPase sites between MCM8/9$^{ATP\gamma S}$ (gold and purple) and MCM8/9$^{ADP}$ (grey, PDB ID 8S91). For both panels, the right-side subunits were aligned, and the movements of the left-side subunits were indicated by black arrows. The ATPγS, ADP and their contacting residues were shown as sticks and the $Mg^{2+}$ was displayed as spheres. **c,** Overall view the ATPγS binding to the CTD ring of MCM8/9 in the MCM8/9$^{ATP\gamma S}$-HROB-DNA complex. The ATPγS molecules were highlighted by their corresponding cryo-EM density maps. The ATPase site at the 9F/8A interface was empty. Comparison of ATPase sites at **d,** 9F/8A, **e,** 8A/9D, **f,** 9D/8B, **g,** 8B/9E, **h,** 9E/8C, and **i,** 8C/9F interfaces in the MCM8/9$^{ATP\gamma S}$-HROB-DNA complex (green and blue) with the corresponding ATPase sites in the MCM8/9$^{ATP\gamma S}$ complex (gold and purple). For all panels, the right-side subunits were aligned and the movements of the left-side subunits were indicated by black arrows.

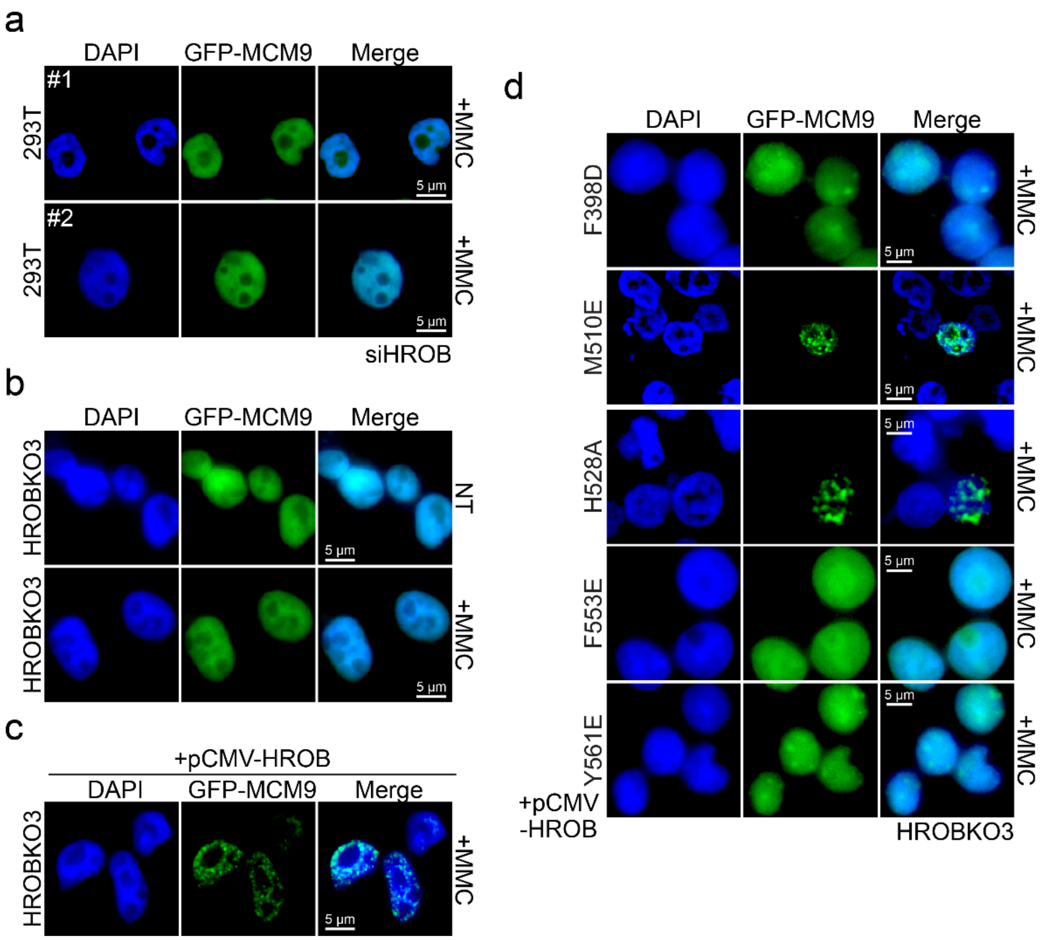


**Supplementary Fig. 10. HROB interaction with MCM8/9 is required for MMC induced nuclear foci formation in human cells.**

MMC induced GFP-MCM9 foci are eliminated with **a,** siRNA knockdown of HROB (#1 and #2 indicate two trials) or in **b,** HROBKO clone 3. **c,** Transfection of WT HROB back into HROBKO cells (clone 3) restores MMC induced GFP-MCM9 foci formation. **d,** Transfection ofpCMV- HROB mutants (F398D, M110E, H528A, F553E, or Y561E) into HROBKO3 to monitor any restoration of MMC induced GFP-MCM9 foci formation.
